# TRAIT: A Toolbox for Rapid Assembly of Integration Templates for CRISPR/Cas- and Integrase-Mediated Genome Engineering

**DOI:** 10.64898/2026.11.17.767390

**Authors:** Jan-Patrick Hetzel, Khaldoon Abdullah, Palomi Schacht, Oleg Timofeev

## Abstract

The generation of genetically engineered mammalian cell models often requires repeated construction of integration donors that differ in their targeting sequences, genetic payloads, or integration strategies. Although modular DNA assembly methods facilitate construct generation, adapting existing expression cassettes to different genome-engineering applications remains labor-intensive. Here, we present TRAIT (Toolbox for Rapid Assembly of Integration Templates), a modular cloning framework that combines direct Golden Gate assembly of targeting elements with Gateway-mediated transfer of genetic payloads. The core vector architecture separates genome-targeting sequences from payload assembly, allowing existing Gateway-compatible constructs to be reused without modification of internal Type IIS restriction sites. We developed vector modules supporting homology-directed repair, homology-independent targeted integration, FrameSelect-compatible donor cleavage, coordinated guide RNA delivery, inducible Cas9 expression, Bxb1-mediated cassette exchange, and conditional transgene expression. An extended MultiSite Gateway workflow enables the assembly of complex integration donors using independently prepared regulatory modules, including components from the Mammalian ToolKit. Functional validation in human cells demonstrated targeted integration, recombinase-mediated cassette exchange, and multiplexed genome editing. We further applied TRAIT to generate human induced pluripotent stem cells carrying functional doxycycline-inducible Cas9 and candidate landing-pad integration constructs. Together, these results establish TRAIT as an interoperable framework for assembling and deploying diverse genome-engineering constructs in mammalian cells.

## Introduction

CRISPR/Cas-based genome engineering has enabled targeted modification of mammalian genomes, including gene disruption, sequence replacement, and integration of genetic payloads into defined genomic loci^1,2^. Targeted integration can be achieved through homology-directed repair (HDR) or homology-independent mechanisms, including nonhomologous end joining (NHEJ) and alternative end joining^3,4^. HDR donor templates contain homology arms flanking the sequence to be integrated, whereas homology-independent targeted integration (HITI) typically employs CRISPR/Cas recognition sequences to promote donor cleavage and integration. Additional approaches, including FrameSelect-based donor cleavage, allow the donor cleavage position to be adjusted to different reading-frame configurations^5,6^. Recombinase-mediated cassette exchange (RMCE) provides a complementary strategy in which genomic landing pads are established before subsequent insertion or exchange of genetic payloads using site-specific recombinases or integrases^7–9^.

Although these approaches differ in their molecular mechanisms, they share a practical challenge: integration templates must be adapted to theintended genomic locus and integration strategy. Repeated integration of different transgenes into the same locus, or deployment of one payload through several integration strategies, often requires construction and validation of multiple donor plasmids that differ only in their targeting elements. This becomes particularly cumbersome when complex expression cassettes have already been assembled and validated, or when open reading frames (ORFs) are available in established plasmid collections. A versatile cloning framework should therefore permit independent modification of genomic targeting sequences and genetic payloads while minimizing unnecessary reconstruction of previously validated DNA modules.

Several modular DNA-assembly technologies address different aspects of this problem. Gateway cloning uses bacteriophage λ-derived site-specific recombination to transfer DNA fragments between compatible entry and destination vectors. Standard Gateway LR recombination enables reuse of existing expression cassettes, while MultiSite Gateway permits assembly of several independently prepared modules within a single destination construct^10,11^. The method is particularly attractive for transferring ORFs and complex expression cassettes without requiring the removal of internal restriction-enzyme recognition sites. However, Gateway assembly depends on compatible recombination sites, retains att-derived junction sequences, and is less suited to unrestricted combinatorial assembly of numerous short DNA fragments.

Golden Gate cloning offers complementary capabilities. This method employs Type IIS restriction endonucleases to generate defined cohesive ends, enabling ordered assembly of multiple fragments in a one-pot restriction–ligation reaction^12^. Depending on the design of the assembly junctions, Golden Gate cloning can preserve seamless connections between DNA elements. Its versatility has led to the development of hierarchical modular cloning systems, including MoClo, EMMA, and the Mammalian ToolKit (MTK), which provide standardized parts for constructing transcriptional units and complex genetic circuits^13–15^. However, direct Golden Gate assembly of pre-existing ORFs or large expression cassettes can be constrained by internal recognition sites for the enzymes employed, potentially requiring sequence domestication or alternative assembly strategies.

The complementary properties of Golden Gate and Gateway cloning have previously been exploited in hybrid assembly systems. Kirchmaier et al. developed Golden GATEway, which combines Golden Gate assembly of predefined DNA components with MultiSite Gateway recombination to generate complex transgenesis constructs^16^ . In this system, genetic elements are prepared in positional Golden Gate entry vectors, assembled into Gateway-compatible intermediate constructs, and subsequently combined through MultiSite LR recombination. This hierarchical organization facilitates reuse of modular components and assembly of sophisticated expression cassettes. Subsequent adaptations have further extended Golden GATEway-derived modular assembly concepts to mammalian expression applications^17^. Nevertheless, genome-integration workflows frequently require custom targeting sequences, such as locus-specific homology arms, donor-cleavage elements, or integrase attachment sites, whose preparation in dedicated positional entry vectors may introduce additional cloning steps.

To address these requirements, we developed TRAIT (Toolbox for Rapid Assembly of Integration Templates), a modular vector framework specifically tailored to mammalian genome engineering. The core Template Cloning Vector (pTCV) contains two flanking Type IIS cloning regions surrounding a Gateway recombination cassette. This arrangement permits direct Golden Gate assembly of appropriately designed targeting fragments into the integration-template backbone, followed by transfer of existing genetic payloads through Gateway LR recombination without reconstructing the flanking elements. An extended MultiSite Gateway architecture further supports independent preparation and combination of regulatory modules, Gateway-compatible payloads, and integration-specific targeting elements. We demonstrate the application of TRAIT to the construction of HDR- and HITI-compatible donors, coordinated guide RNA delivery, inducible genome-editing systems, Bxb1-compatible landing pads and cassette-exchange donors, and conditional transgene-expression constructs. Functional experiments in HEK293-derived cells and human induced pluripotent stem cells establish the utility of this modular framework across several mammalian genome-engineering workflows.

## Results

### Generation of the Template Cloning Vector

Construction of CRISPR/Cas-mediated integration templates requires the incorporation of strategy-specific targeting sequences, such as homology arms for homology-directed repair (HDR) or short recognition sequences for homology-independent targeted integration (HITI). These elements can be readily generated by PCR or synthesized as double-stranded oligonucleotides, making them suitable for Golden Gate assembly. In contrast, complex genetic payloads may contain internal Type IIS restriction sites that complicate direct Golden Gate cloning or require sequence domestication.

To facilitate the reuse of existing payloads in different genome-integration strategies, we developed the Template Cloning Vector (pTCV), the core component of the TRAIT toolbox. The vector contains two tandem Type IIS cloning regions flanking an attR1–ccdB–CmR–attR2 Gateway recombination cassette (Figure 1A). This architecture separates the assembly of integration-specific targeting elements from the introduction of genetic payloads. Targeting sequences are first assembled directly into the corresponding flanking sites by Golden Gate cloning, after which a payload from a compatible Gateway entry vector is transferred by LR recombination (Figure 1B).

**Figure 1.**
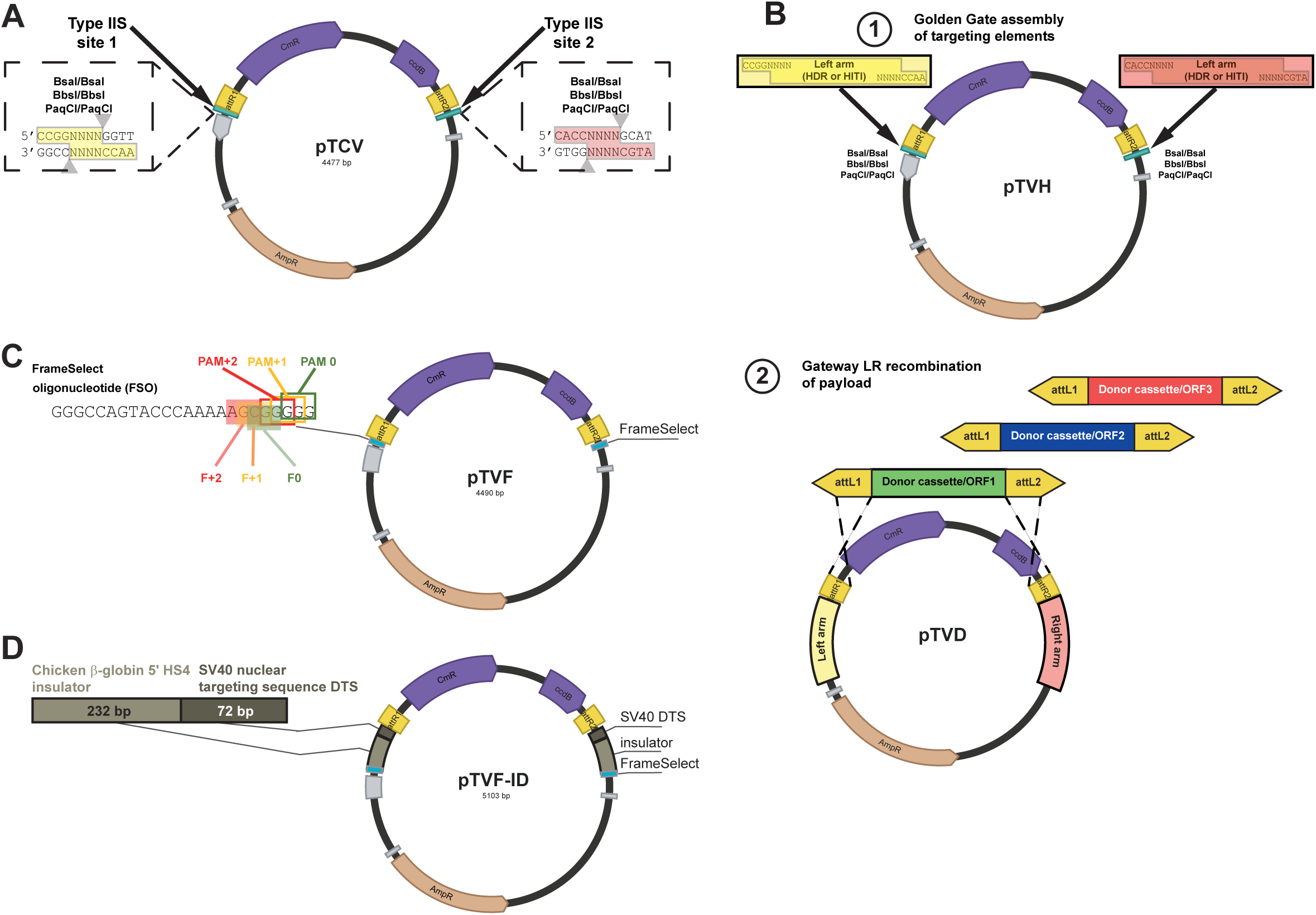
Design and architecture of the TRAIT cloning system for generation of mammalian genome-integration templates. (A) Schematic representation of the Template Cloning Vector (pTCV), the core component of TRAIT. A Gateway attR1–ccdB–chloramphenicol-resistance–attR2 cassette is flanked by two tandem Type IIS restriction-site regions, enabling independent Golden Gate assembly of sequences positioned upstream and downstream of the genetic payload. The available pTCV variants contain BsaI/BsaI, BbsI/BbsI, PaqCI/PaqCI, or BsaI/BbsI cloning-site combinations. (B) Two-step workflow for construction of genome-integration donors. In the first step, integration-specific targeting elements, such as homology arms or short targeting sequences, are assembled into the flanking pTCV cloning sites by Golden Gate assembly. In the second step, a genetic payload is transferred from a compatible Gateway entry vector into the prepared targeting backbone by LR recombination. The targeting backbone can subsequently be reused with alternative payloads without reconstructing the flanking elements. (C) Design of the FrameSelect-compatible pTVF vector. FrameSelect recognition sequences flank the Gateway cassette and permit cleavage at defined nucleotide offsets using corresponding FrameSelector sgRNAs, enabling adjustment of donor cleavage positions for the intended reading-frame configuration during HITI-mediated integration. (D) Design of pTVF-ID, incorporating flanking chicken β-globin HS4-derived core insulator elements (Ins) and the SV40 DNA nuclear targeting sequence (DTS) in addition to the FrameSelect modules. These elements were included to support subsequent targeted-integration applications.

Unlike hierarchical Golden GATEway workflows that employ positional Golden Gate entry vectors^16^, this arrangement allows appropriately designed PCR fragments, synthetic oligonucleotides, and compatible modular DNA parts to be assembled directly into the pTCV backbone. The resulting targeting vector can subsequently be reused with different Gateway payloads without reconstructing the targeting elements. The principal architectural differences between TRAIT and established Golden Gate, Gateway, and Golden GATEway workflows are summarized in Table 1.

**Table 1.**
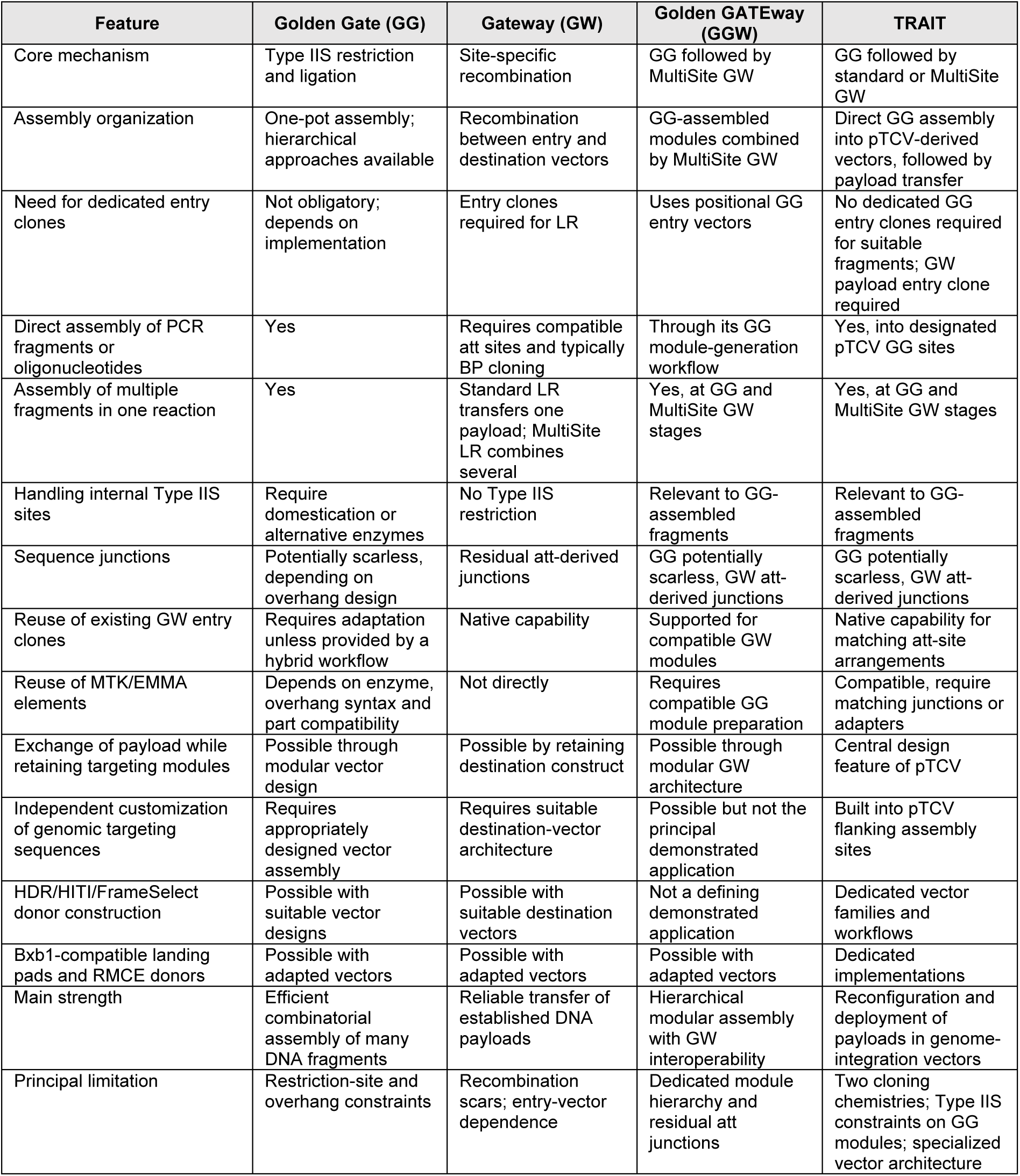
Comparison of Golden Gate, Gateway, Golden GATEway and TRAIT for modular DNA assembly and construction of genome-engineering vectors.

To increase the flexibility of the system, we generated four pTCV variants carrying BsaI/BsaI, BbsI/BbsI, PaqCI/PaqCI, or BsaI/BbsI cloning tandems. We next adapted pTCV for HITI-based in-frame integration by incorporating FrameSelector (FS) sequences. In this design, the donor template is flanked by short sequences recognized by FrameSelector gRNAs, which cleave at one of three adjacent nucleotide positions corresponding to the three possible coding frames^5,6^. This design allows the donor cleavage position to be adjusted to the intended reading-frame offset, facilitating in-frame integration without reconstructing the complete donor cassette for each target configuration. However, the final reading frame remains dependent on the sequence of the integration junction. Using this strategy, we generated a pTVF vector containing FS sequences that can be cleaved at positions 0, +1, or +2 (Figure 1C).

Because expression of integrated transgenes can be affected by surrounding chromatin, we also generated a modified donor-template vector containing flanking insulator sequences to reduce position effects and support stable transgene expression 22. In addition, we included an SV40 DNA nuclear targeting sequence (DTS), which has been reported to enhance nuclear import of plasmid DNA^18^. The resulting plasmid was named pTVF-ID (Figure 1D).

We tested our pTVH and pTVF-ID donor vectors for Cas9-mediated gene targeting. Long homology arms for targeting the human safe-harbor locus *hROSA26/THUMPDS3-AS1* were introduced into pTCV to generate the pTVH-hR26 donor backbone. A payload for integration – a GFP-puromycin resistance cassette, was then inserted by LR recombination into this vector, as well as into the pTVF-ID vector to generate donor templates pTVH-hR26-GFP-puro and pTVF-ID-GFP-puro for HDR- and HITI-mediated integration, respectively. HEK293 cells were co-transfected with a Cas9 expressing plasmid, the corresponding GFP-puro donor template and the respective sgRNA vectors (Figure S1A). We analyzed GFP, RFP657 and mCherry expression (latter are the markers of sgRNA-vectors) by flow cytometry to assess transfection efficiency (Figure S1B), and selected transfected cells with puromycin. Transfection with HITI-compatible vectors produced at least as many puromycin-resistant colonies as the HDR-based approach under the conditions tested, suggesting comparable or higher recovery of stable integrants. Similar results were obtained upon targeting of the GFP-puro HDR or HITI donors into *AAVS1* safe harbor locus ( Figure S1C). Because antibiotic selection does not distinguish targeted from ectopic integration, these results reflect the recovery of resistant colonies rather than the efficiency of precise genome targeting.

### Using TRAIT for generation of cells with targeted integration of inducible CRISPR/Cas9

Retroviral and lentiviral vectors are frequently used to generate isogenic cell lines with stable inducible expression of Cas9. However, because these vectors integrate randomly into the genome and may be affected by LTR silencing, they are not always optimal for generating well-defined engineered cell models. We therefore sought to generate donor templates for CRISPR/Cas9-mediated targeted integration of inducible Cas9 cassettes into genomic safe-harbor loci.

Using the pTCV architecture, we generated a family of iCG (inducible Cas9) donor plasmids compatible with HDR-driven targeted integration or HITI (Table S1). These constructs contain tetracycline-inducible Cas9 co-expressed with EGFP, together with a reverse tetracycline transactivator and a neomycin-resistance marker (Figure 2A,B). The iCGC construct additionally contains Cre recombinase fused to a mutated estrogen receptor ligand-binding domain (CreERT2). In this system, CreERT2 is retained in the cytoplasm until treatment with 4-hydroxytamoxifen (4-OHT), which enables nuclear translocation and Cre-mediated recombination between loxP sites, thereby supporting conditional gene expression (Figure 2A). Combining inducible Cas9 with inducible Cre activity increases the versatility of the system by enabling both CRISPR/Cas-mediated editing and recombinase-controlled genetic switches in the same cell line. Inducible EGFP expression from these constructs was first tested after transient transfection into HEK293 cells and treatment with doxycycline (Figure S2A).

**Figure 2.**
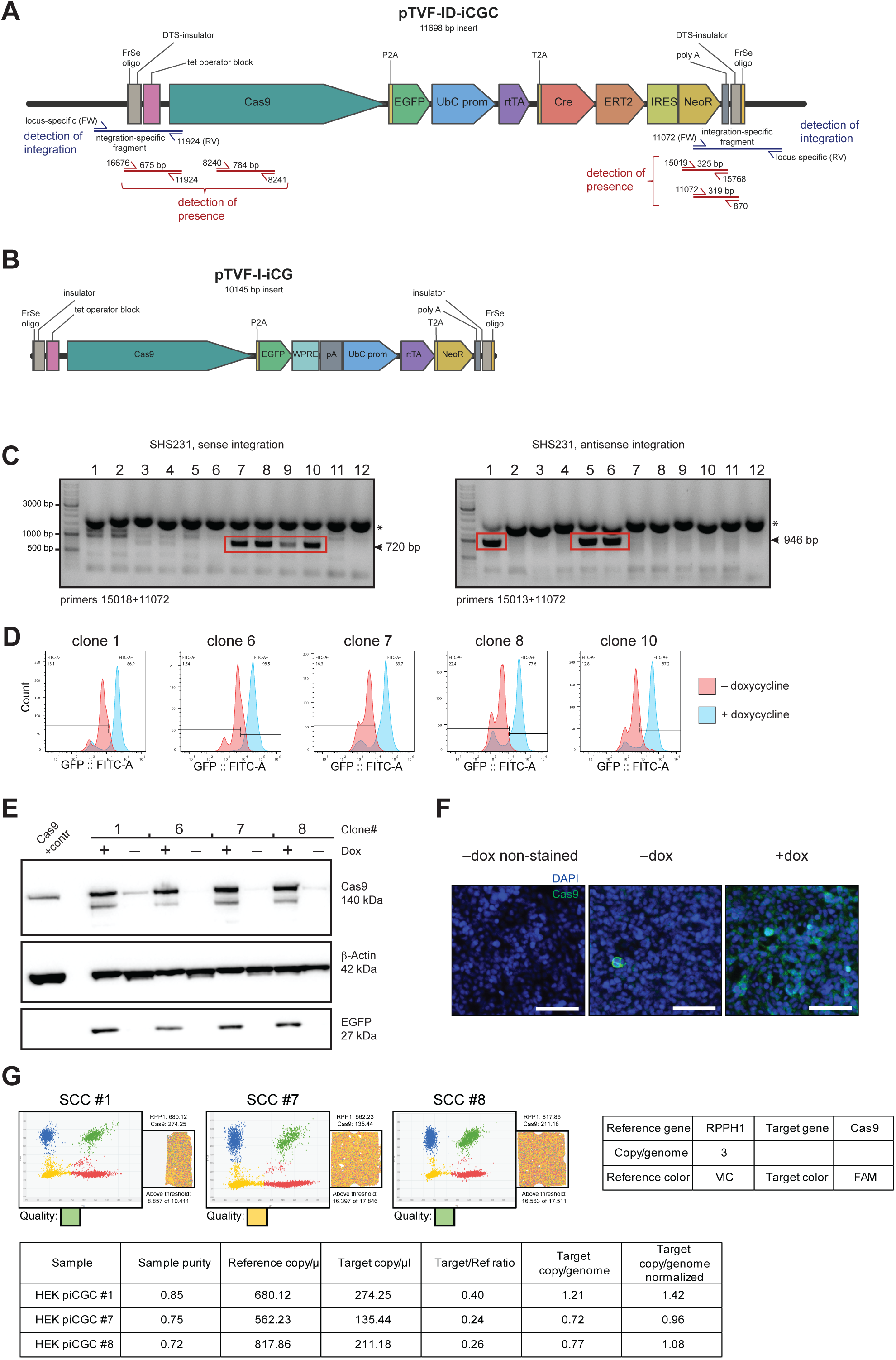
**Generation and characterization of HEK293 cells carrying targeted integrations of inducible Cas9 expression cassettes**. (A, B) Schematic representation of the TRAIT-derived donor constructs pTVF-ID-iCGC (A) and pTVF-I-iCG (B). Both contain a tetracycline-responsive Cas9 expression cassette linked to an EGFP reporter, a constitutively expressed reverse tetracycline transactivator (rtTA), and a neomycin-resistance marker. The iCGC construct additionally encodes Cre^ERT2^, enabling recombinase-mediated genetic modification. FrameSelect sequences and flanking insulator/DTS elements support construction of HITI-compatible integration donors. Combinations of primers used for PCR to confirm integration (shown in blue, pairs of locus-specific and construct-specific primers) or presence of the construct (shown in red, pairs of construct-specific primers) are shown in (A). (C) Integration-specific PCR analysis of neomycin-resistant HEK293 single-cell-derived clones following HITI-mediated integration of the iCGC cassette into the *SHS231* locus. PCR assays were designed to detect integration in either orientation, as indicated. The expression cassette is functional independently of its genomic orientation. Red boxes and arrowheads indicate the specific PCR products, asterisk – unspecific bands. Primers used for PCR are shown below. (D) Representative flow-cytometry histograms showing doxycycline-inducible EGFP expression in five selected HEK293-iCGC clones. Cells were treated with 1 µg/mL doxycycline for 48 h or maintained without doxycycline. The indicated percentages represent the fractions of EGFP-negative and EGFP-positive cells. (E) Western blot analysis of Cas9 and EGFP expression in the indicated clones following treatment with 1 µg/mL doxycycline for 48 h (+) or in untreated controls (−). β-Actin served as a loading control. (F) Representative immunofluorescence images showing doxycycline-dependent Cas9 expression in HEK293-iCGC cells. Cas9 immunostaining is shown in green and nuclei are counterstained with DAPI (blue). Cells were treated with 1 µg/mL doxycycline for 48 h. Scale bar, 125 µm. (G) Digital PCR analysis of Cas9 transgene copy number in three selected HEK293-iCGC clones using *RPPH1* as the reference locus. The analysis employed a three-copy *RPPH1* reference model based on the reported HEK293 karyotype.

We next used the iCGC construct for targeted integration into the *SHS231* locus, which has been characterized as a reliable genomic safe harbor^19^. HEK293 cells were co-transfected with piCGC together with two SGL40C.EFS.RFP657 plasmids expressing sgRNAs targeting the *SHS231* locus and the flanking sequences in the integration template. Transfected cells were selected with neomycin and resistant single-cell clones (SSC) were isolated and screened by integration-specific PCR (Figure 2C). SSC with confirmed targeted integration were further analyzed for doxycycline-inducible Cas9-EGFP expression by flow cytometry, immunofluorescence, and western blotting (Figure 2D,E,F). To determine transgene copy number, three PCR-positive clones were analyzed by dPCR. Digital PCR detected one copy of the integrated transgene in each of the three analyzed clones (Figure 2G).

Because CreERT2-based systems can show inducer-independent background activity, we next assessed Cre activity using a pTCV-based INTERSECT Con/Fon fluorescent reporter^20,21^. This reporter expresses mScarlet only when both Cre and Flpo recombinases are active in the same cell (Figure S2B). Parental HEK293 cells and stable inducible SSC HEK293-iCGC were transfected with the Con/Fon reporter alone or together with a Flpo-expression plasmid. In parental cells, the reporter remained silent despite Flpo expression, confirming that Flpo alone was insufficient to activate mScarlet expression. In contrast, a substantial proportion of HEK293-iCGC cells co-transfected with the Con/Fon reporter and Flpo showed high mScarlet expression even without 4-OHT, and treatment with 4-OHT only slightly increased the number of mScarlet-positive cells (Figure S2C,D). These results demonstrated functional Cre-mediated recombination in HEK293-iCGC cells, but also revealed substantial 4-OHT-independent recombinase activity. This can potentially be explained by activity of the CreERT2 cytoplasmic pool that caused recombination of the reporter plasmid after transfection, because the Cre recombinase domain is inherently active regardless of the presence of 4-OHT^22^.

### Generation and validation of TRAIT sgRNA and donor-template vectors

In addition to construction of donor-templates, TRAIT can be used to support coordinated delivery of sgRNAs required for HDR- and HITI-mediated genome engineering. The pTCV architecture allows PCR-amplified U6-sgRNA cassettes to be inserted into one or both flanking cloning sites, generating vectors that express single or multiple sgRNAs. For HITI using pTVF donor templates, two sgRNA activities are required: a FrameSelector sgRNA cleaves the FS sequence in the donor plasmid to release the integration template (Figure 1C), whereas another sgRNA (or paired sgRNAs) target the genomic integration site. Based on pTCV architecture, we designed two families of gRNA vectors to be used with the pTVF-ID donor vectors for HITI – pTVF-g and pTVF-go, that enable co-delivery of target-specific sgRNAs together with the corresponding FrameSelector sgRNA from a single plasmid.

The first pTVF-g family contains a fixed FrameSelector sgRNA expression cassette inserted into either the left or right cloning site of pTCV. The second Type IIS tandem, BsaI or BbsI, respectively, remains available for insertion of target-specific U6-sgRNA expression cassettes. This design is useful for subcloning validated sgRNAs from existing plasmids or for assembling multiple U6-sgRNA cassettes into one block (Figure 3A). We then used these constructs to generate the second pTVF-go family of vectors that contain on one flank a U6 promoter and Type IIS cloning sites for Golden Gate insertion of sgRNA oligonucleotides, and the FrameSelector sgRNA block positioned on the other flank of the ccdB cassette (Figure 3B). In parallel, we produced a set of pENTR donor plasmids carrying different combinations of fluorescent and antibiotic-resistance markers, which can be transferred by Gateway LR recombination into pTVF, pTVF-g/go and other pTCV-based vectors (Figure 3C, Table S1). Together, these plasmids expand TRAIT from donor-template assembly to coordinated delivery of payloads and sgRNA modules.

**Figure 3.**
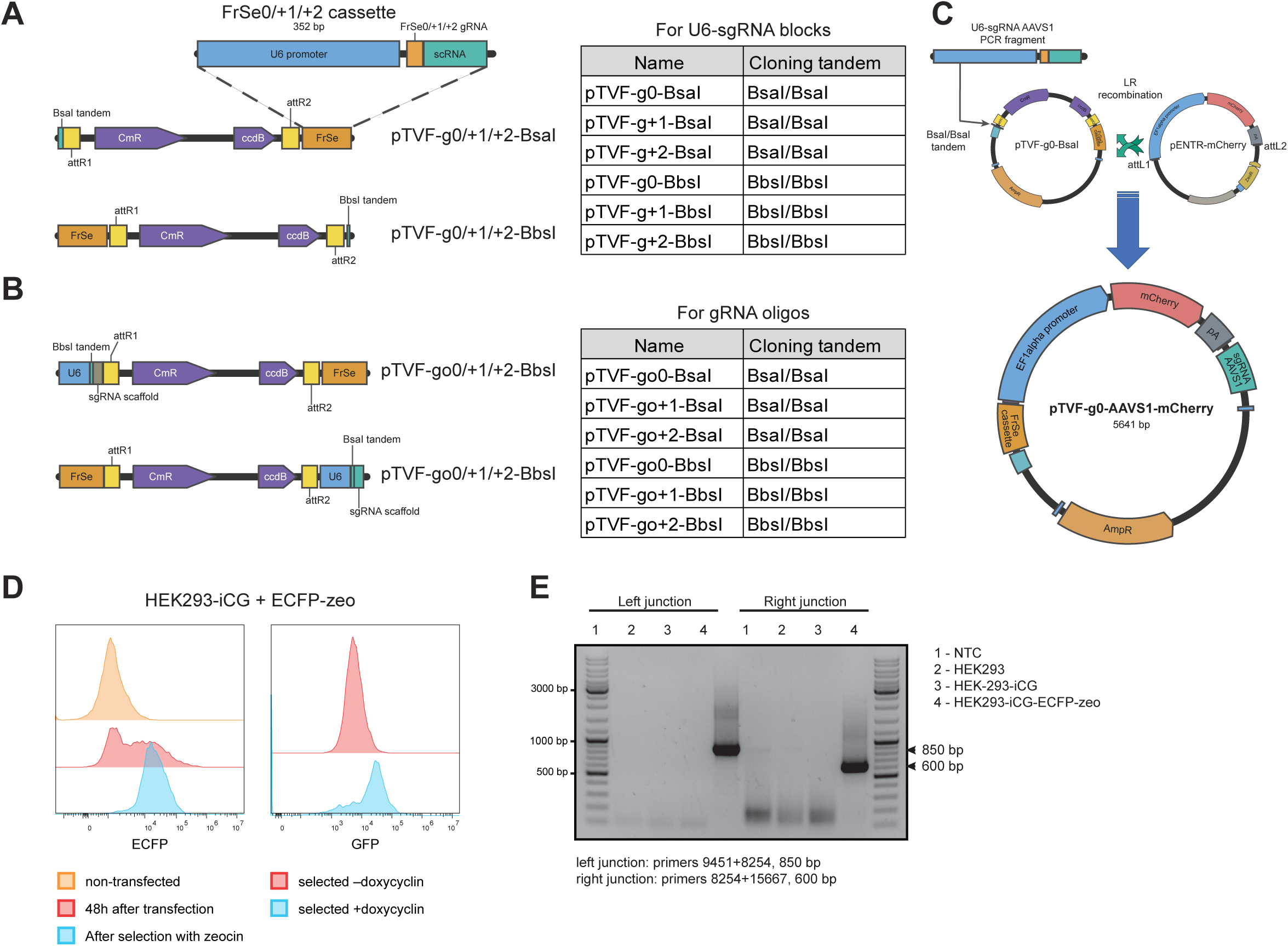
**Design and application of TRAIT sgRNA-expression vectors for coordinated donor cleavage and genomic targeting**. (A) Schematic representation of the pTVF-g vector family, which combines a fixed U6-driven FrameSelector sgRNA expression cassette with an available Type IIS cloning region for insertion of target-specific sgRNA-expression modules. FrameSelector sgRNAs recognizing cleavage positions 0, +1, or +2 are positioned in one BsaI or BbsI cloning tandem, while the second tandem remains available for cloning. The six vector configurations and their corresponding cloning enzymes are listed. (B) Schematic representation of the pTVF-go vector family, designed for direct Golden Gate insertion of target-specific sgRNA oligonucleotides. These vectors contain a U6 promoter and sgRNA scaffold adjacent to the corresponding Type IIS cloning sites, together with a separate FrameSelector sgRNA-expression cassette. Six configurations are provided, combining three FrameSelect cleavage positions with BsaI or BbsI cloning sites. (C) Representative assembly workflow for a combined sgRNA-expression and fluorescent-reporter construct. A PCR-amplified U6-sgRNA cassette targeting *AAVS1* locus is inserted into pTVF-g0-BsaI by Golden Gate assembly. Subsequent Gateway LR recombination with pENTR-mCherry generates pTVF-g0-AAVS1-mCherry, which combines the genomic-targeting sgRNA, the FrameSelector sgRNA and an mCherry expression cassette in a single plasmid. (D) Flow-cytometric analysis of HEK293-iCGC cells after cotransfection with the TRAIT HITI donor pTVF-ID-ECFP-zeo and the combined sgRNA vector pTVF-g0-AAVS1-mCherry, followed by doxycycline induction and Zeocin selection. ECFP and mCherry fluorescence are used to monitor the donor and sgRNA-vector-associated populations, respectively. (E) Integration-specific PCR analysis of genomic DNA from the selected ECFP-positive population, designed to detect targeted donor integration at the *AAVS1* locus. Nested PCRs with indicated primers were performed using as template the purified PCR fragments obtained from gDNA with primer pairs 15678/17007 and 17008/15667 for the left and right junctions, respectively (Figure S3B).

We next evaluated the combined functionality of inducible Cas9, TRAIT sgRNA-expression vectors, and targeted-integration donors in HEK293-iCGC cells. The cells were co-transfected with the pTVF-ID-ECFP-zeo donor template and the pTVF-g0-AAVS1-mCherry vector, carrying FrameSelector 0 sgRNA and a validated sgRNA specific for the *AAVS1* locus. After transfection, Cas9 expression was induced by doxycycline for 48 h. Cells with stable integration were selected with Zeocin and analyzed by FACS, which confirmed a strong enrichment of ECFP population after selection, and showed that selected cells retain inducible GFP expression (Figure 3D). Genomic DNA from the pool of selected cells was analyzed by integration-specific PCR which detected donor–genome junction products consistent with targeted integration at *AAVS1* (Figure 3E, Figure S3A,B). Sanger sequencing of PCR-amplified fragments confirmed the presence of correctly targeted *AAVS1* junctions in the selected ECFP-positive population ( Figure S3C,D), supporting the combined functionality of the system.

### TRAIT enables construction of landing-pad and inducible sgRNA modules

CRISPR/Cas-mediated targeted integration, particularly of large donor templates, is often inefficient and requires extensive selection and validation of modified clones^3^. Combining CRISPR/Cas-mediated insertion of genomic landing pads with recombinase-mediated cassette exchange (RMCE) overcomes some of these limitations. Large serine integrases mediate site-specific integration of attB-flanked donor cassettes into attP-containing landing pads^23–25^. This strategy was proven efficient for precise integration of large payloads and assembly of complex expression cassettes in living cells^7–9,26^.

We used the pTVF-ID architecture to generate landing-pad templates compatible with HDR- or HITI-mediated integration and generated two modified vectors: pTVF-ID-PaqCI and pTVF-ID-BbsI-PaqCI. The first plasmid contains two PaqCI tandem cloning sites flanking the ccdB cassette, enabling cloning of attP oligos for integrase recombination using Golden Gate assembly (Figure S4A). The second modification carries two additional Bbs tandem cloning sites next to the FS oligo, which provides sites for insertion of homology arms, making the construct suitable for CRISPR-Cas9 mediated targeting via HDR and HITI ( Figure S4B). Different fluorescent and antibiotic-resistance markers are placed into landing pad vectors by Gateway LR recombination and used for selection of targeted cells (Figure 1B). Restricting the initial targeted-integration step to a comparatively small landing-pad cassette was intended to simplify establishment of recipient cell lines before subsequent integration of larger payloads via Bxb1-mediated recombination of attB-flanked donor cassettes. These donor plasmids can be easily generated from the same pTVF-ID-PaqCI and pTVF-ID-BbsI-PaqCI vectors using Golden Gate insertion of respective attB oligos, followed by LR recombination with an entry plasmid carrying a cargo of interest.

We next used this landing pad concept to design an inducible sgRNA-expression system compatible with RMCE. Temporal regulation of genome editing is often desirable and can be achieved with inducible Cas9 systems, such as the iCG and iCGC constructs described above. However, in cell lines that already constitutively express Cas9, delivery of conventional sgRNA cassettes driven by Pol III promoters, such as U6 or U3, leads to immediate formation of Cas9-sgRNA complexes. Temporal control of sgRNA production can instead be achieved by expressing sgRNAs from inducible Pol II promoters. Because RNA-cleaving systems can process sgRNAs post-transcriptionally from a Pol II transcript^27^, we used the ribozyme-gRNA-ribozyme (RGR) strategy, in which hammerhead (HH) and hepatitis delta virus (HDV) ribozymes flank the 5′ and 3′ ends of the sgRNA, respectively, and release the mature sgRNA by self-cleavage^28^.

The system comprises two components: a CRISPR/Cas-targetable landing pad and an attB-flanked donor cassette for Bxb1 recombinase-mediated cassette exchange (RMCE). The landing-pad vector, pTVF-ID-gRNA-GFP-puro-LP-attP(Bxb1), contains a tetracycline-inducible promoter, a GFP-puromycin resistance cassette flanked by two orthogonal Bxb1 attP sites, and a promoterless zeocin-resistance gene (Figure 4A). Following targeted integration, GFP expression and puromycin resistance enable selection and identification of landing-pad-positive cells. The corresponding RMCE donor carries two compatible orthogonal Bxb1 attB sites flanking an ECFP reporter, one or more ribozyme-flanked sgRNA (RGR-sgRNA) cassettes, and a downstream PGK promoter (Figure 4B). Bxb1-mediated cassette exchange replaces the GFP-puromycin cassette with the donor payload, thereby reconstituting a tetracycline-inducible sgRNA-expression module and placing the PGK promoter in front of the zeocin-resistance marker (Figure 4C). In the resulting configuration, the tetracycline-inducible promoter drives co-transcription of ECFP and the RGR-sgRNA cassette(s). A 110-bp MALAT1-derived triple-helix element positioned downstream of ECFP was incorporated to promote poly(A)-independent stabilization of the upstream transcript 35. The downstream PGK promoter drives constitutive expression of the zeocin-resistance gene, enabling selection of cells carrying the exchanged cassette (Figure 4C).

**Figure 4.**
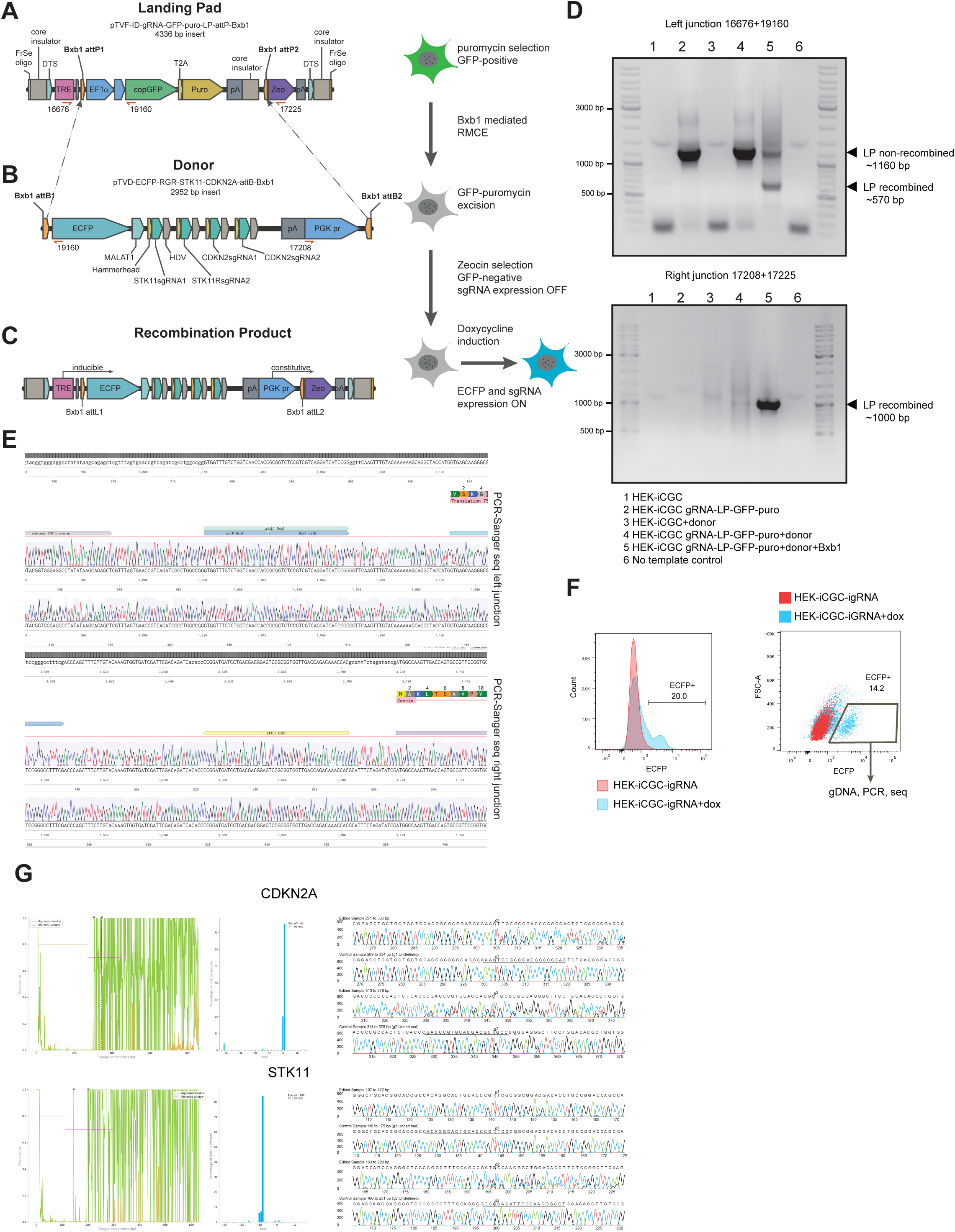
**TRAIT-mediated construction of a Bxb1 landing-pad system for inducible expression of multiple CRISPR sgRNAs**. (A) Schematic representation of the pTVF-ID-gRNA-GFP-puro-LP-attP(Bxb1) landing-pad construct. The cassette contains a tetracycline-responsive promoter (TRE), a constitutively expressed GFP–puromycin-resistance selection module flanked by orthogonal Bxb1 attP sites, and a promoterless zeocin-resistance marker. FrameSelect recognition sites, insulator elements, and SV40 DNA nuclear targeting sequences (DTS) support construction of a targeted-integration donor. The corresponding primer-binding positions used for junction PCR are indicated. (B) Structure of the pTVD-ECFP-RGR-STK11-CDKN2A-attB(Bxb1) donor cassette containing an ECFP reporter, four ribozyme-flanked sgRNA expression modules targeting *STK11* and *CDKN2A*, and a downstream PGK promoter. Two orthogonal Bxb1 attB sites flank the exchangeable cassette. The MALAT1-derived stabilization element and hammerhead and hepatitis delta virus (HDV) ribozyme sequences are indicated. (A-C) Schematic representation of the Bxb1-mediated cassette-exchange workflow. Recombination between compatible attP and attB sites replaces the GFP–puromycin cassette with the ECFP–RGR-sgRNA donor and positions the PGK promoter upstream of the zeocin-resistance marker. Zeocin selection enriches cells carrying the exchanged cassette. Subsequent doxycycline induction activates ECFP and RGR-sgRNA expression. (D) Integration-junction PCR analysis of parental HEK293-iCGC cells, landing-pad-containing cells, and cells transfected with the RGR-sgRNA donor with or without Bxb1 integrase. Primer pairs 16676/19160 and 17208/17225 were used to examine the left and right recombination junctions, respectively. The sample identities and size of fragments are indicated alongside the gels. (E) Representative Sanger sequencing chromatograms of PCR products spanning the left and right recombination junctions, confirming the expected donor–landing-pad junction sequences following Bxb1-mediated cassette exchange. (F) Flow-cytometric analysis of ECFP expression in HEK293-iCGC cells carrying the exchanged inducible RGR-sgRNA cassette, cultured with or without doxycycline. The histogram and scatter plot show ECFP-positive fractions in two experiments. The ECFP-positive population was enriched by fluorescence-activated cell sorting for subsequent genomic DNA isolation and editing analysis. (G) Sanger sequencing and ICE (Inference of CRISPR Edits) analysis of the *STK11* and *CDKN2A* target regions in doxycycline-treated, ECFP-positive cells carrying the integrated inducible RGR-sgRNA cassette. Representative sequencing profiles from edited and control samples are shown alongside ICE analysis outputs. Editing frequencies were estimated from the Sanger sequencing data in the enriched cell population.

To validate the sgRNAs for the inducible expression system, we designed two pairs of sgRNAs targeting exon 5 of *STK11* and exon 2 of *CDKN2A*, respectively. Each sgRNA pair was cloned into the pSpCas9(BB)-2A-Puro vector and tested in HEK293 cells. Four days after transfection, genomic DNA was isolated and the targeted regions were amplified by PCR. The detection of shorter amplicons was consistent with sgRNA-mediated deletions at both loci (Figure S4C).

To test the inducible sgRNA system, we generated HEK293-iCGC cells with stable integration of the pTVF-ID-gRNA-GFP-puro-LP-attP (Bxb1) landing pad (HEK293-iCGC-gRNA-LP). The four sgRNAs, whose combined activity was confirmed by deletion PCR, were subsequently assembled into the pTVD-ECFP-RGR-STK11-CDKN2A-attB(Bxb1) donor plasmid, and used together with a Bxb1 integrase-expression plasmid for co-transfection of HEK293-iCGC-gRNA-LP cells. Successful cassette exchange was verified by junction PCR and Sanger sequencing, confirming replacement of the GFP-puromycin cassette with the RGR-sgRNA donor cassette (Figure 4D,E). Following zeocin selection, resistant cells were treated with doxycycline for four days to induce Cas9 and RGR-sgRNA expression, and ECFP-positive cells were enriched by FACS (Figure 4F). Genomic DNA from sorted cells was used for PCR amplification of the *STK11* and *CDKN2A* targeted fragments ( Figure S4D) and Sanger sequencing. Sanger sequencing followed by ICE analysis revealed extensive editing at the *STK11* and *CDKN2A* target regions in the doxycycline-treated, ECFP-positive population (Figure 4G). Together, these results demonstrate that TRAIT-assembled landing-pad and RGR-sgRNA modules support Bxb1-mediated cassette exchange and functional expression of multiple sgRNAs in human cells.

### TRAIT tools for assembly of complex constructs by MultiSite Gateway

Although the original pTCV architecture facilitates the transfer of payloads from existing pENTR/pDONR vectors into integration templates, assembling more complex constructs requires additional regulatory elements to be cloned together with genome-targeting sequences. This reduces flexibility when combining different payloads, regulatory modules, and targeting strategies. To overcome this limitation, we extended TRAIT with MultiSite Gateway cloning, which enables the assembly of multiple DNA fragments through orthogonal att recombination sites in a single LR reaction^10,11^.

We generated two complementary pTCV-derived vectors: pTCV-attL4-attR1-attR2-attL3 and pTCVKanA-attR4-attR3 (Figure 5A,B). The first retains the original pTCV architecture but incorporates attL4 and attL3 sites flanking the two BsaI tandem cloning regions, allowing additional regulatory elements, such as promoters, tags, poly(A) signals, or expression cassettes, to be introduced by Golden Gate assembly. The second serves as the MultiSite Gateway destination backbone and contains an attR4/attR3-flanked ccdB cassette, PaqCI tandem cloning sites for inserting genome-targeting elements, and a kanamycin-resistance marker for selection of the final construct (Figure 5B).

**Figure 5.**
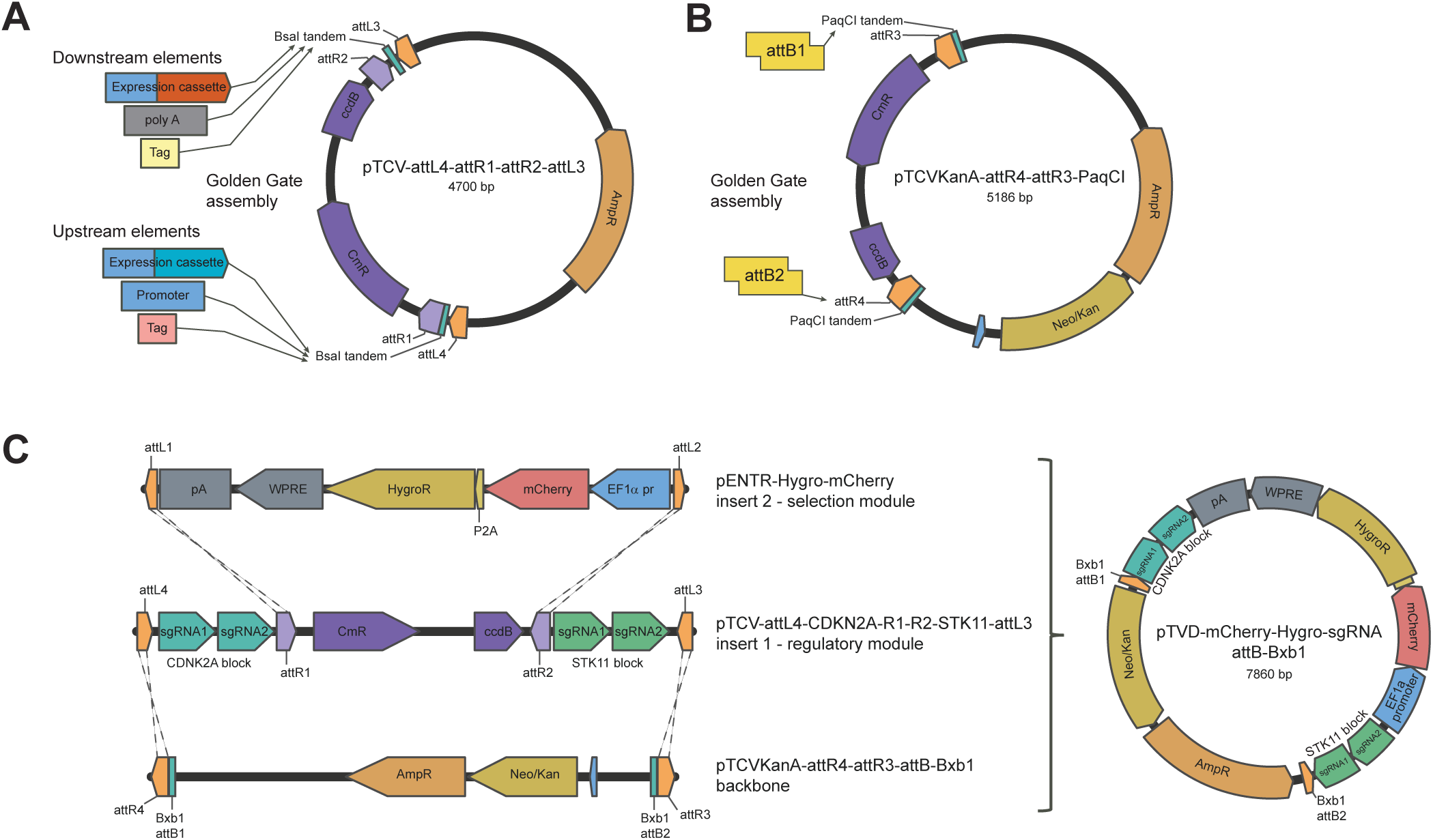
Extension of TRAIT for hierarchical assembly of complex genome-integration constructs using MultiSite Gateway recombination. **n**(A) Structure of pTCV-attL4-attR1-attR2-attL3, a Gateway-compatible vector containing two flanking BsaI tandem cloning regions for Golden Gate assembly of additional regulatory or functional elements. The attL4 and attL3 recombination sites permit incorporation of these elements during subsequent MultiSite Gateway assembly. (B) Structure of pTCVKanA-attR4-attR3-PaqCI, which serves as the destination backbone for MultiSite Gateway assembly. Two PaqCI tandem cloning regions enable insertion of genome-targeting elements, including orthogonal Bxb1 attB attachment sites, while the kanamycin-resistance marker permits selection of the resulting destination-derived plasmids. (C) Representative three-component assembly of a complex Bxb1-compatible donor vector. An intermediate containing four U6-sgRNA expression cassettes targeting *CDKN2A* and *STK11* was combined with a Gateway entry construct encoding fluorescent and antibiotic-resistance markers and a destination backbone containing orthogonal Bxb1 attB sites. MultiSite Gateway LR recombination generated pTVD-mCherry-hygro-sgRNA-CDKN2A-STK11-attB(Bxb1). The final plasmid was validated by diagnostic restriction digestion and Sanger sequencing of selected regions and assembly junctions. The resulting construct is designed for integration into landing pads carrying compatible Bxb1 attP sites.

The resulting workflow separates the assembly of targeting sequences from the construction of complex expression modules (Figure 5C). For example, Bxb1 attB sites for recombinase-mediated cassette exchange can be introduced into pTCVKanA-attR4-attR3 through PaqCI-based Golden Gate cloning, while regulatory elements are assembled independently into the BsaI cloning regions of pTCV-attL4-attR1-attR2-attL3. These two constructs are subsequently combined with a payload from a standard pENTR/pDONR vector in a single MultiSite Gateway LR reaction. The resulting integration template is recovered by kanamycin selection in ccdB-sensitive bacteria, which counterselects unrecombined destination plasmids.

To validate this assembly strategy, we generated pTCV-attL4-sgRNA-CDKN2A-attR1-attR2-sgRNA-STK11-attL3, carrying two U6-sgRNA expression cassettes targeting CDKN2A in the left BsaI cloning region and two targeting *STK11* in the right region. This construct was combined with pTCVKanA-attR4-attR3-attB(Bxb1) and pENTR-mCherry-hygro, providing fluorescent and antibiotic-resistance markers, in a MultiSite Gateway LR reaction (Figure 5C). The resulting plasmid, pTVD-mCherry-hygro-sgRNA-CDKN2A-STK11-attB(Bxb1), was verified by construct-specific restriction digestion analysis and Sanger sequencing. The assembled donor contains four sgRNA expression cassettes and is designed for integration into genomic landing pads carrying compatible Bxb1 attP sites. These results demonstrate the feasibility of combining independently assembled regulatory and functional modules into complex integration templates using the extended TRAIT architecture.

### TRAIT tools for conditional transgene expression and compatibility with MTK

Conditional transgene expression is commonly achieved through Cre- or Flpo-mediated excision of transcriptional STOP cassettes. Using TRAIT, we generated a Conditional Expression Cloning Cassette (CECC) for targeted integration and recombinase-dependent activation of transgenes (Figure 6A). The CECC contains an EF1α promoter separated from a multiple cloning site (MCS) by a loxP-flanked STOP cassette encoding EYFP and blasticidin resistance. Downstream of the MCS, a T2A ribosome-skipping sequence links transgene expression to an RFP657 fluorescent reporter. Before Cre-mediated excision, EYFP expression and blasticidin resistance enable identification and selection of cells carrying the cassette, whereas excision of the STOP sequence permits expression of the transgene and RFP657 reporter (Figure 6A).

**Figure 6.**
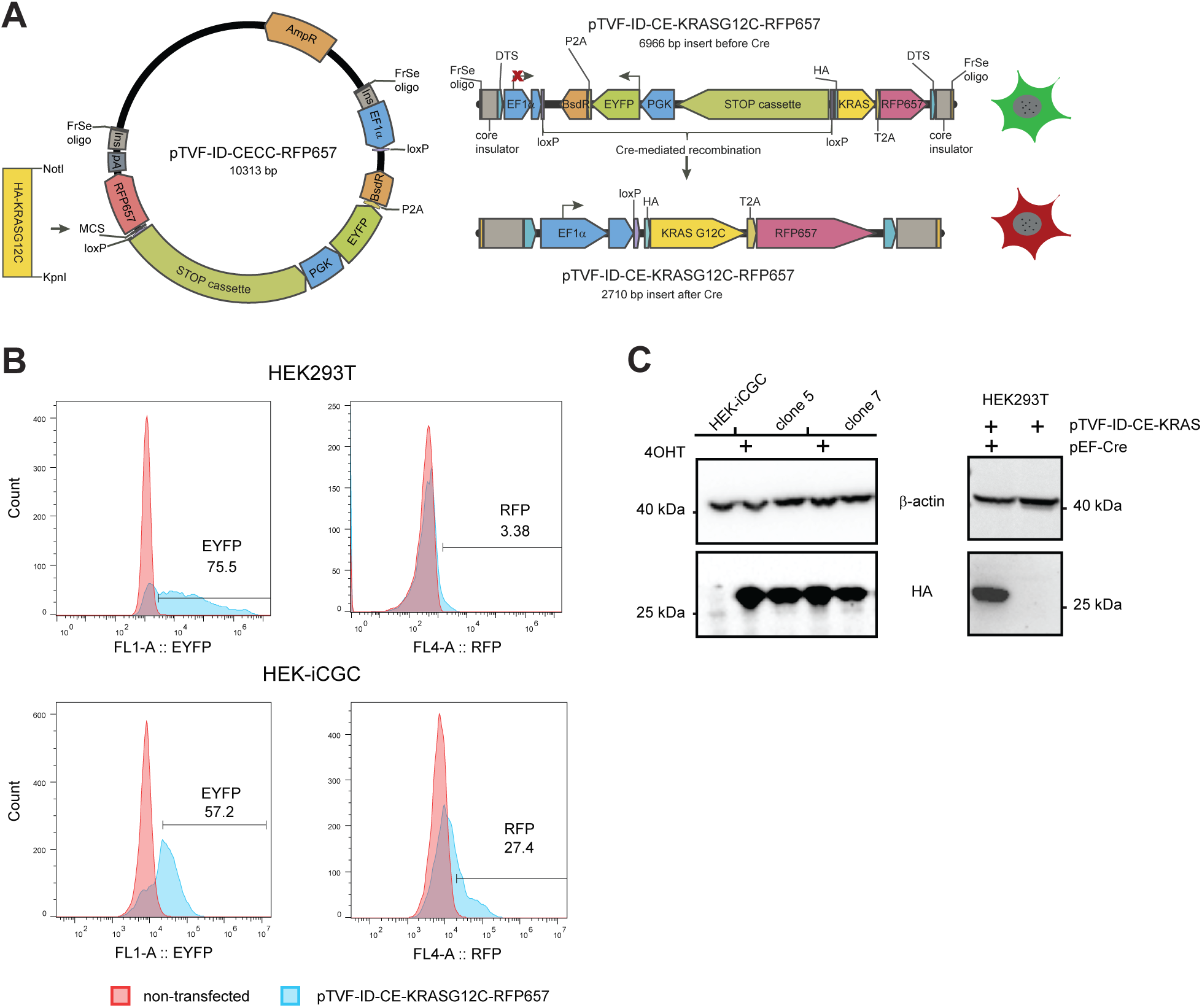
Design and functional evaluation of a TRAIT-derived conditional transgene-expression cassette. (A) Schematic representation of the conditional-expression cloning cassette (CECC) and its Cre-mediated activation. The cassette contains an EF1α promoter separated from the HA-tagged KRAS^G12C transgene and downstream T2A-RFP657 reporter by a loxP-flanked transcriptional STOP cassette encoding EYFP and blasticidin resistance. Before recombination, the cassette permits EYFP expression and antibiotic selection, whereas Cre-mediated excision of the STOP region enables expression of HA-KRAS^G12C and RFP657. (B) Representative flow-cytometry histograms showing EYFP and RFP657 expression in parental HEK293 and HEK293-iCGC cells transfected with the conditional KRAS^G12C^ construct. Reporter-positive fractions are indicated. (C) Western blot analysis of HA-KRAS^G12C^ expression in selected HEK293-iCGC-derived clones and parental HEK293 cells transfected with the conditional donor. HA-tagged KRAS was detected in the analyzed HEK293-iCGC clones and following cotransfection of parental HEK293 cells with a Cre-expression plasmid. β-Actin served as the protein-loading control. The experimental conditions, including 4-OHT and Cre-expression plasmid treatment, are indicated above the corresponding lanes.

To evaluate CECC functionality, we inserted HA-tagged KRAS^G12C^ into pTVF-ID-CECC and used the resulting pTVF-ID-CE-HA-KRAS^G12C^ construct for *AAVS1*-targeted integration in HEK293-iCGC cells or transient transfection of parental HEK293 cells. Following blasticidin selection, both transfections resulted in enrichment of EYFP-positive cell populations (Figure 6B). Parental HEK293 cells showed no detectable RFP657 fluorescence, whereas HEK293-iCGC cells expressed RFP657 and HA-KRAS^G12C^ even without 4-hydroxytamoxifen (4-OHT) treatment (Figure 6B; Figure S5A). This observation is consistent with the ligand-independent CreERT2 activity detected in the Con/Fon reporter assay ( Figure S2C,D) and suggests that spontaneous STOP-cassette excision may occur in the pool of transfected donor plasmid, potentially before stable integration, and while EYFP and resistance marker expression is provided by non-recombined plasmids co-expressed in the same cell. Consistently, all analyzed blasticidin-resistant HEK293-iCGC-KRAS^G12C^ clones expressed exogenous HA-KRAS^G12C^, as confirmed by western blotting (Figure 6C). In contrast, HA-KRAS^G12C^ expression in parental HEK293 cells required co-transfection with a Cre-expression plasmid (Figure 6C).

To demonstrate the compatibility of TRAIT with existing modular cloning libraries, we extended the conditional-expression platform to incorporate additional oncogenic payloads and regulatory elements. Gateway entry constructs were generated for several oncogenes, including KRAS^G12D^, EGFR^L858R^, and PIK3CA^E545K^ (Table S1). Because these complex cassettes contain internal recognition sites for Type IIS restriction enzymes used in Golden Gate assembly, their direct use in such workflows would require sequence modification or alternative assembly strategies. TRAIT circumvents this limitation by enabling the transfer of intact payloads through Gateway recombination. As a proof of principle, we adapted the conditional KRAS^G12C^ construct for Bxb1-mediated integration and incorporated a WPRE–poly(A) module downstream of the RFP657 reporter. WPRE and poly(A) elements from the Mammalian ToolKit (MTK) were assembled with compatible adapter oligonucleotides into pTCV-attL4-attR1-attR2-attL3 by Golden Gate cloning. The assembled module was additionally flanked by PaqCI recognition sites to facilitate its reuse in subsequent cloning reactions ( Figure S5B). The resulting intermediate was then combined with the existing pENTR-CE-HA-KRAS^G12C^-RFP657 cassette and the pTCVKanA-attR4-attR3-attB(Bxb1) destination vector in a MultiSite Gateway LR reaction, generating pTVD-CE-HA-KRAS^G12C^-attB(Bxb1) ( Figure S5C). Correct assembly was confirmed by diagnostic restriction digestion and Sanger sequencing. Thus, TRAIT enabled the combination of MTK-derived regulatory elements, a preassembled conditional-expression cassette, and a recombinase-targeting backbone without sequence modification of the original transgene cassette, demonstrating interoperability between distinct modular cloning strategies.

### Using TRAIT for generation of genetically modified iPSCs

After validating the TRAIT system in HEK293 cells, we next asked whether it could be used to generate genetically modified human induced pluripotent stem cells (iPSCs). iPSCs provide a particularly useful model for genome engineering because, unlike tissue-specific adult stem cells, they can give rise to multiple cell lineages. This expands the utility of genetically modified iPSCs for the generation of diverse organoid models and for the assembly of complex multicellular systems that more closely mimic native tissues^29^.

We transfected two iPSC lines, WTSIi004A and SCVI-15, with an iCG donor template for HITI along with validated sgRNA constructs targeting two safe harbor loci - *hROSA26* and *SHS231*. Transfected cells were selected with neomycin, and gDNA from the resulting resistant cells was used for analysis with landing pad and junction-specific PCRs, which confirmed presence of targeted cells in the obtained pools (Figure 7A and Figure S5A). Despite PCR evidence consistent with targeted integration, detectable doxycycline-induced GFP expression varied between the tested cell-line and genomic-locus combinations: while in WTSIi004A with iCG integration into *hROSA26* locus approximately 30% of cells were GFP-positive after doxycycline treatment (Figure 7B), the SCVI-15 and WTSIi004A pools and clones with confirmed integration in *SHS231* locus showed no inducible GFP expression (Figure S5B). From 34 GFP-positive SSC obtained for WTSIi004A with iCG targeted into *hROSA26* locus, five clones with the highest levels of GFP induction were selected for further characterization (Figure 7C). After confirming doxycycline-inducible Cas9 expression by immunofluorescence (Figure 7D, Figure S5C), we analyzed the selected clones for expression of pluripotency markers to ensure that generated cells have not lost pluripotency during selection. Flow cytometry showed that clone #28 has expression of TRA-1-60-R, TRA-1-81, SSEA-3 and SSEA-4 similar to the parental cell line, and we therefore selected this clone for further experiments (Figure 7E).

**Figure 7.**
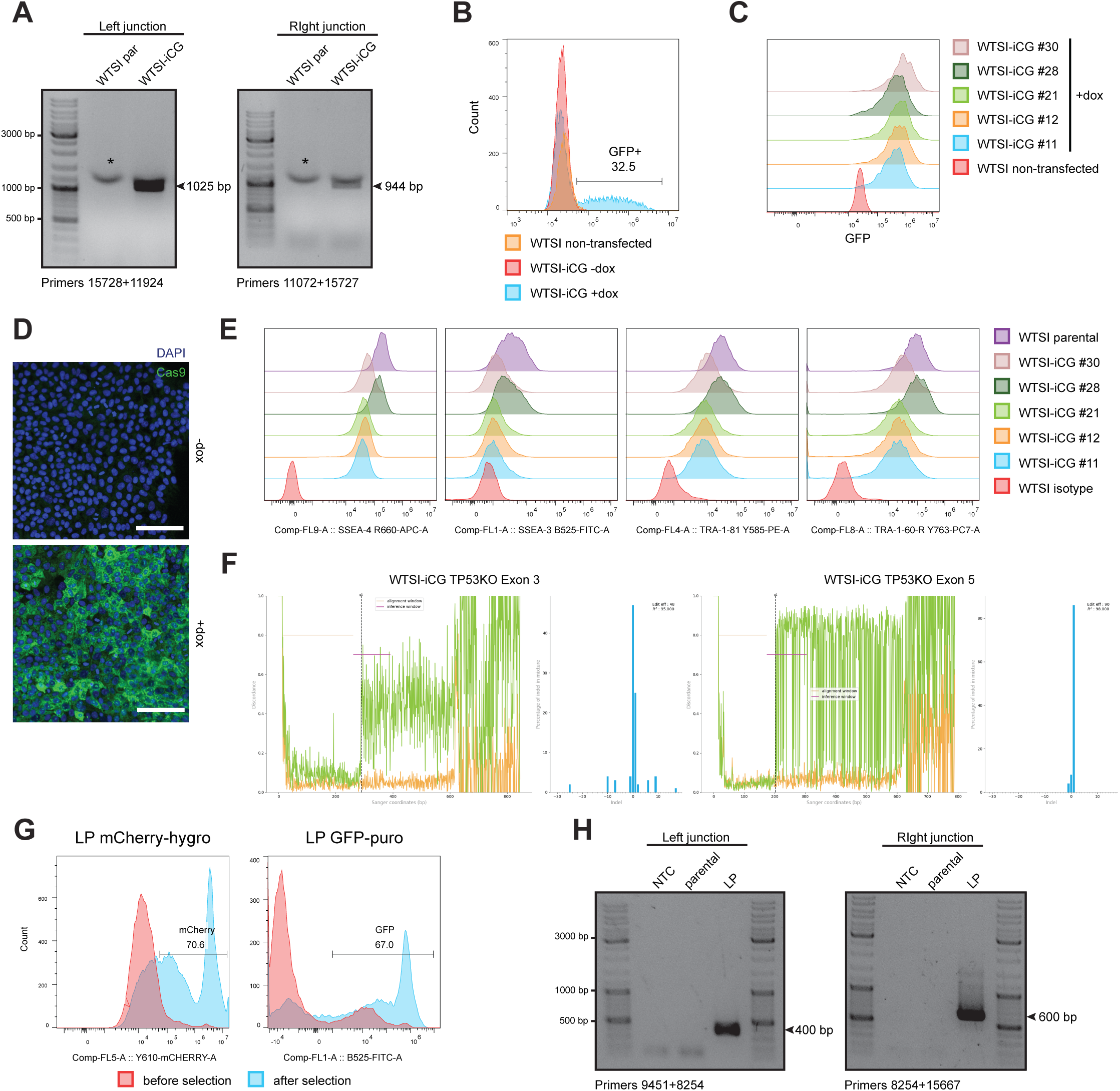
Generation and characterization of human iPSC lines carrying TRAIT-derived inducible Cas9 and landing-pad constructs. (A) Integration-specific PCR screening of neomycin-resistant human iPSC populations following HITI-mediated targeting of inducible Cas9 (iCG) donor constructs to the *hROSA26/THUMPDS3-AS1* locus for the WTSIi004A (WTSI) iPSC line. The indicated amplicons correspond to the expected integration-specific products, asterisk indicates unspecific bands. (B) Flow-cytometric analysis of doxycycline-responsive EGFP expression in WTSI-iCG cells following targeting of the iCG cassette to *hROSA26*. Fluorescence profiles are shown for parental WTSI cells and engineered cells cultured with or without doxycycline. (C) Flow-cytometry histograms showing EGFP expression in selected single-cell-derived WTSI-iCG clones following doxycycline treatment. A parental WTSI population is included as a control. (D) Representative immunofluorescence images showing Cas9 expression in WTSI-iCG cells cultured with or without doxycycline (1 µg/mL, 48 h). Cas9 immunostaining is shown in green, and nuclei are counterstained with DAPI (blue). Scale bar, 125 µm. (E) Flow-cytometric assessment of pluripotency-associated surface markers SSEA-4, SSEA-3, TRA-1-81, and TRA-1-60-R in the indicated WTSI-iCG clones, compared with parental WTSI cells and isotype controls. (F) Sanger sequencing and ICE (Inference of CRISPR Edits) analysis of the *TP53* exon 3 and exon 5 target regions in WTSI-iCG clone #28 pool following sgRNA transfection, doxycycline induction of Cas9, and Nutlin-3A selection. ICE analysis was performed on the Nutlin-3A-selected cell population. (G) Flow-cytometry histograms showing reporter enrichment in WTSI-iCG populations following introduction of TRAIT-derived landing-pad GFP-puro and mCherry-hygro donor constructs after antibiotic selection. The left panel shows mCherry fluorescence following selection with hygromycin (125 µg/mL); the right panel shows GFP fluorescence following selection with puromycin (1 µg/mL). Fluorescence profiles before and after selection are shown. (H) Integration-specific PCR analysis of genomic DNA from the WTSI-iCG#28 (parental) and the selected puromycin-resistant, GFP-positive population (LP), designed to detect targeted donor integration of the GFP-puro landing pad at the *AAVS1* locus. NTC – no template control.

To assess functionality of the inducible Cas9 in the generated WTSI-iCG clone #28 we transfected cells with two validated sgRNAs targeting exons 3 and 5 of human *TP53*, and Cas9 expression was induced by continuous doxycycline treatment for four days. Next, cells were selected with increasing concentrations (1-5 μM) of Mdm2 inhibitor Nutlin-3A, which induces stress-independent activation of p53 (ref). Multiple clones grew after selection of transfected cells, whereas non-transfected cells died in a few days of treatment. To confirm *TP53* editing, we used genomic DNA isolated from pool of selected cells and amplified the targeted region by PCR. Sanger sequencing followed by ICE analysis identified editing in both *TP53* targeted exons in the Nutlin-3A-selected population, consistent with enrichment of cells carrying disruptive *TP53* alterations. (Figure 7F and Figure S5D,E).

We also generated additional iPSC lines suitable for future RMCE-based engineering. For this purpose, different landing pads were targeted into the *hROSA26* and *AAVS1* safe-harbor loci of SCVI-15 and WTSI-iCG cells, respectively. The polyclonal antibiotic-resistant, reporter-positive iPSC populations were analyzed by PCR, which confirmed presence of the landing pads and provided evidence for targeted integration (Figure 7G,H and Figure S5F). Thus, we obtained candidate iPS cell lines for subsequent molecular validation and RMCE-based engineering.

## Discussion

The development of TRAIT was motivated by a recurrent challenge in mammalian genome engineering: the construction of integration donors that share a common targeting configuration but differ in their genetic payloads, or conversely, donors that carry the same payload but require different targeting elements. TRAIT addresses this problem through a modular vector architecture in which genome-targeting sequences and genetic payloads can be assembled and modified independently. Type IIS Golden Gate cloning enables the introduction of locus- or application-specific elements into flanking cloning regions, whereas Gateway recombination facilitates the transfer of previously assembled and validated expression cassettes. This organization allows targeting modules to be reused with different payloads and reduces the need to reconstruct complete donor plasmids for each genome-engineering application. The resulting toolbox encompasses constructs for CRISPR/Cas-mediated integration, controlled expression of genome-editing components, and integrase-mediated cassette exchange.

TRAIT builds on established DNA-assembly technologies rather than introducing a new recombination or ligation mechanism. In particular, the Golden GATEway system described by Kirchmaier et al. (2013) demonstrated the utility of combining Golden Gate assembly with MultiSite Gateway recombination to construct complex genetic elements. TRAIT differs primarily in its organization around mammalian genome-integration workflows. While Golden GATEway uses positional Golden Gate entry vectors to prepare modules for subsequent Gateway-mediated assembly, the basic TRAIT architecture permits appropriately designed targeting fragments to be introduced directly into the flanking cloning regions of a Gateway-compatible destination vector. Existing Gateway entry clones can then provide payload sequences without requiring removal of internal Type IIS recognition sites. This arrangement is particularly suited to the recurrent exchange of transgenes, reporters, or selection cassettes within established targeting configurations. Moreover, the extended MultiSite Gateway workflow allows independently prepared targeting and regulatory modules to be combined with existing payloads. The incorporation of MTK-derived WPRE and polyadenylation elements illustrates how components from established modular cloning libraries can be incorporated into TRAIT constructs. Nevertheless, the choice between TRAIT and other assembly platforms will depend on the intended application, available plasmid resources, and desired assembly complexity.

The development of donor vectors for HDR-dependent and homology-independent targeted integration (HITI) demonstrates how a common assembly architecture can accommodate different genomic targeting mechanisms. HDR donors require homology arms, whereas HITI strategies employ donor-cleavage sequences that enable integration through end-joining repair pathways. The FrameSelect-compatible TRAIT derivatives additionally allow adjustment of the donor cleavage position for applications requiring integration in a defined reading frame. In our experiments, HITI-based approaches supported the recovery of antibiotic-resistant cells following delivery of the corresponding donor constructs. However, the number of resistant colonies should not be interpreted as the frequency of correctly targeted integration, because selection can also recover cells carrying unintended or rearranged insertions. Accordingly, the principal advance of these vectors is their modular design and adaptability to different targeting strategies rather than a demonstrated improvement in genomic integration efficiency. Their utility was further supported by the generation of iPSC populations with integration-associated PCR products and, in selected clones, functional inducible Cas9 expression.

A particularly useful extension of TRAIT is its compatibility with landing-pad strategies and recombinase-mediated cassette exchange (RMCE). Direct CRISPR/Cas-mediated insertion of large expression constructs can require repeated optimization, selection, and clonal validation whenever the payload is changed. Landing-pad systems offer an alternative in which a genomic docking site is established once and subsequently reused for the introduction of different payloads through site-specific recombination. We generated Bxb1-compatible landing-pad and donor vectors and demonstrated cassette exchange in HEK293-derived cells using an inducible reporter and guide RNA expression cassette. Junction-specific PCR and Sanger sequencing supported the expected recombination products in selected cell populations, while fluorescence analysis provided an additional readout of donor expression. These experiments establish the functional compatibility of the TRAIT donor architecture with Bxb1-mediated exchange.

The inducible ribozyme–guide RNA–ribozyme (RGR) module further extends the platform by enabling conditional production of guide RNAs from a polymerase II transcript. This approach complements conventional U6-driven sgRNA expression and may be useful when guide RNA production must be coordinated with other inducible genetic components. In the tested system, a tetracycline-responsive promoter controlled expression of an ECFP-associated RGR cassette containing multiple guide RNA sequences targeting *STK11* and *CDKN2A* genes. Following cassette exchange, induction and enrichment of ECFP-positive cells, molecular analysis identified editing at the intended target regions. Thus, the experiment demonstrates the feasibility of combining landing-pad-mediated donor exchange, inducible guide RNA expression, and Cas9-mediated genome editing within a single workflow.

TRAIT also supports the construction of conditional transgene-expression systems. The Conditional Expression Cloning Cassette (CECC) incorporates a loxP-flanked transcriptional STOP cassette between a constitutive promoter and the transgene of interest, allowing selection and detection of cells carrying the unrecombined construct before Cre-mediated activation. Using HA-tagged KRAS^G12C^ and a linked RFP657 reporter, we generated and characterized conditional oncogene-expression constructs. The modular architecture further enabled incorporation of the same expression cassette into a Bxb1-compatible donor together with independently prepared regulatory elements. These examples illustrate how established payloads can be transferred between conventional and integrase-compatible vector configurations without reconstructing the entire expression cassette.

At the same time, experiments with inducible Cre^ERT^^2^ revealed an important limitation of conditional recombinase systems: appreciable reporter activation occurred in the absence of 4-hydroxytamoxifen. The observed ligand-independent activity indicates that nominally inducible recombinases can exhibit substantial background activity under particular experimental conditions. These findings emphasize the importance of experimentally assessing both induced activity and background recombination when implementing conditional-expression modules.

Application of TRAIT to human induced pluripotent stem cells (iPSCs) extends the experimental scope of the toolbox beyond readily transfectable HEK293-derived cells. We introduced inducible Cas9 constructs into two human iPSC lines and obtained integration-associated PCR products at the targeted genomic loci. However, doxycycline-responsive reporter expression differed considerably among the tested targeting configurations and cell populations. Robust inducible expression was obtained in selected WTSIi004A-derived clones following targeting of the *hROSA26* locus. One of these clones was used for Cas9-mediated editing of *TP53* gene. We also recovered antibiotic-resistant, fluorescent reporter-positive iPSC populations following delivery of TRAIT landing-pad constructs. Although these results support the applicability of the cloning framework to human iPSCs, targeted integration and functional cassette exchange remain to be established for the candidate iPSC landing-pad populations. The observed variability between integration-associated PCR detection and inducible transgene expression additionally highlights the importance of screening individual targeting configurations and clones for functional activity.

Several technical limitations should be considered when applying TRAIT. First, modular donor assembly simplifies plasmid construction but does not eliminate uncertainties associated with genomic integration. HITI and other end-joining-based approaches can produce indels at integration junctions, donor rearrangements, or unintended insertions, while HDR-based targeting is subject to cell-type- and locus-dependent constraints. Antibiotic resistance, fluorescence, and internal donor PCR are useful screening readouts but cannot substitute for direct validation of both genomic integration junctions and, where necessary, the integrity and copy number of the inserted cassette. Second, the use of Gateway recombination introduces defined att-derived junction sequences, which must be considered when designing protein fusions or regulatory elements that require precise sequence continuity. Third, complex multi-fragment assembly and the introduction of large genetic payloads may require construct-specific optimization; neither the assembly performance nor the genomic integration efficiency of TRAIT vectors has been systematically benchmarked against alternative platforms. Finally, conditional expression and editing systems require functional validation in each cellular context, as illustrated by Cre^ERT2^ background activity and variability in inducible Cas9 expression among iPSC populations.

Future development of TRAIT could expand the range of compatible genomic loci, integrase systems, and regulatory modules. In particular, establishing and validating stable recipient cell lines with characterized landing pads could facilitate recurrent integration of large expression constructs, variant libraries, or synthetic gene circuits without repeating the initial CRISPR/Cas targeting procedure. Additional improvements may include strategies for reducing background recombinase activity, optimizing inducible guide RNA expression, and incorporating alternative chromatin-regulatory elements or donor-delivery formats. Because TRAIT uses established cloning chemistries and is compatible with existing Gateway entry constructs and selected Golden Gate modules, it may be particularly useful for laboratories seeking to adapt existing molecular resources to diverse mammalian genome-engineering applications. The availability of the TRAIT plasmid collection through Addgene is intended to facilitate adoption, further characterization, and community-driven development of the platform.

In summary, TRAIT provides an integration-oriented framework that connects established modular DNA-assembly methods with CRISPR/Cas- and integrase-based mammalian genome engineering. By allowing targeting elements and genetic payloads to be prepared and exchanged independently, it supports the construction of diverse donor templates and associated genetic-control modules. Its application to targeted integration, cassette exchange, inducible genome editing, conditional expression, and human iPSC engineering demonstrates the versatility of the approach while identifying opportunities for further optimization and validation.

## Methods

### Generation of TRAIT plasmids

A complete list of TRAIT plasmids is provided in Table S1, and the sequences of all oligonucleotides used for the cloning procedures described below are provided in Table S2. Unless otherwise indicated, restriction enzymes and reagents for restriction–ligation and Golden Gate assembly were obtained from New England Biolabs (NEB). Plasmid DNA was prepared using the ZymoPURE II Plasmid Mini- or Midiprep Kit (Zymo Research). Constructs generated by Gibson assembly were verified by diagnostic restriction digestion and Sanger sequencing across the complete assembled region. Golden Gate- and Gateway-derived constructs were verified by diagnostic restriction digestion and Sanger sequencing of assembly junctions and selected complete inserts.

The pTCV cloning vector was generated from a PCR-amplified ccdB cassette flanked by Gateway attR1 and attR2 sites. The cassette was amplified from pAd/PL-DEST (Thermo Fisher Scientific) using primers containing KpnI and tandem BsaI sites at the 5′ end and XbaI and tandem BbsI sites at the 3′ end. The resulting 1,793-bp fragment was cloned into pCR-Blunt II-TOPO using the Zero Blunt TOPO PCR Cloning Kit (Invitrogen), sequence-verified using M13 forward and reverse primers, and subsequently transferred into pUC57 by KpnI/XbaI restriction and ligation. Endogenous BsaI sites in the ampicillin-resistance cassette and ccdB sequence were removed by site-directed mutagenesis using silent G-to-A and C-to-G substitutions, respectively. The resulting pTCV vector contained tandem BsaI and BbsI cloning sites flanking the Gateway destination cassette. Derivatives containing BsaI, BbsI, or PaqCI cloning tandems were generated by Golden Gate insertion of appropriate double-stranded oligonucleotides.

To generate FrameSelect-compatible pTVF vectors, FrameSelect oligonucleotides derived from the system described by Schmid-Burgk et al. (2016) were extended with overhangs compatible with the corresponding pTCV cloning sites and introduced by Golden Gate assembly. For generation of the insulated pTVF-ID derivatives, insulator sequences were PCR-amplified from pHR700PA-1 37 . Primers were designed to introduce the appropriate FrameSelect sequences and self-removing BsaI sites for Golden Gate assembly. The resulting PCR products, together with double-stranded oligonucleotides encoding DNA targeting sequences (DTS), were assembled into the corresponding pTCV derivative to generate pTVF-ID vectors.

To generate the pTVF-g0, pTVF-g+1, and pTVF-g+2 family, FrameSelector sgRNAs corresponding to the three FrameSelect reading frames were first cloned into pSpCas9(BB)-2A-Puro (PX459) v2.0 (Addgene). The complete U6–sgRNA expression cassettes were subsequently PCR-amplified and inserted by Golden Gate assembly into the left BsaI or right BbsI tandem sites of pTCV, generating BsaI- and BbsI-based pTVF-g derivatives. For generation of the corresponding pTVF-go vectors, U6-driven guide-RNA cloning cassettes containing BsaI or BbsI sites were inserted into the available cloning tandem of the respective pTVF-g vector. This produced six pTVF-go derivatives corresponding to the three FrameSelect phases and the two restriction-enzyme configurations. Where indicated, an EF1α-driven nuclear mCherry cassette was introduced by Gateway LR recombination using a domesticated pENTR-EF1α-mCherry-NLS-poly(A) entry clone lacking internal BsaI and BbsI sites.

The inducible Cas9 constructs iCG and iCGC were assembled using NEBuilder HiFi DNA Assembly Master Mix (NEB). Fragments encoding the Tet-responsive promoter, FLAG-Cas9-NLS-P2A cassette, EGFP, UbC-rtTA-Cre^ERT2^, and IRES-Neo-poly(A) sequences were amplified from existing plasmids using primers containing approximately 30-bp overlaps and assembled into PCR-linearized pUC57. Gateway attB1 and attB2 sites were incorporated into the outer amplification primers to permit subsequent BP recombination. The iCG variant was generated from the iCGC cassette using Gibson Assembly. The resulting pUC57-iCG and pUC57-iCGC constructs were transferred into pDONR-zeo by Gateway BP recombination and subsequently into the appropriate TRAIT integration-vector backbone by Gateway LR recombination to generate the corresponding FrameSelect-compatible HITI donor templates.

For HDR-mediated integration, pTVH donor backbones carrying homology arms for the human *THUMPDS3-AS1/ROSA26* (*hROSA26*) or *PPP1R12C-201/AAVS1* safe-harbor loci were generated by PCR amplification of the corresponding homology arms using primers containing terminal BsaI or BbsI sites. These fragments were inserted into the left and right pTCV cloning sites by Golden Gate assembly. The resulting pTVH backbones were subsequently combined with Gateway entry clones by LR recombination to generate the final HDR donor templates.

Bxb1 landing-pad vectors were generated from pTVF-ID derivatives. First, FS-insulator-DTS modules extended with PaqCI tandems or PaqCI/BbsI double tandems were introduced into pTCV-BsaI to generate pTVF-ID-PaqCI and pTVF-ID-BbsI-PaqCI intermediates, respectively. Orthogonal Bxb1 attP sequences containing GT or GA central dinucleotides were subsequently introduced into the PaqCI cloning sites by Golden Gate assembly, producing pTVF-ID-attP(Bxb1) and pTVF-ID-BbsI-attP(Bxb1), respectively. Gateway LR recombination with entry clones encoding fluorescent and selectable markers generated final landing-pad constructs, including pTVF-ID-GFP-puro-attP(Bxb1). Corresponding pTVD donor backbones were generated by insertion of the matched orthogonal Bxb1 attB sites into pTCV derivatives followed by Gateway LR recombination with the required payload entry clone.

The pTVF-ID-gRNA-GFP-puro-LP-attP(Bxb1) landing-pad plasmid was assembled in three stages. First, pTVF-ID-gRNA-LP was generated by Gibson assembly of a plasmid backbone containing the ampicillin-resistance cassette, an FS-insulator-DTS-Tet-promoter module and poly(A) sequence amplified from a pTVF-ID-iCG derivative, a ccdB cassette flanked by BbsI cloning tandems amplified from pTCV-BbsI, and a Zeocin-resistance cassette amplified from pDONR-zeo. Orthogonal Bxb1 attP sequences were then introduced into the available BbsI cloning sites by Golden Gate assembly to generate pTVF-ID-gRNA-LP-attP(Bxb1). Finally, Gateway LR recombination with pENTR-GFP-puro generated pTVF-ID-gRNA-GFP-puro-LP-attP(Bxb1).

The corresponding multiplexed RGR donor was generated from an ECFP-RGR entry cassette. Fragments encoding ECFP, the MALAT1-derived RGR scaffold, multiple cloning region, self-cleaving ribozyme elements and the associated expression cassette were assembled into pUC57 by Gibson assembly. Gateway attB1 and attB2 sites were introduced with the outer assembly primers, permitting BP recombination with pDONR-zeo to generate pENTR-ECFP-RGR. This entry clone was transferred by LR recombination into a pTCV-derived Bxb1-attB donor backbone. Individual sgRNA modules containing BbsI-compatible ends, a PCR adaptor, a hammerhead ribozyme, and prevalidated sgRNAs targeting *STK11* or *CDKN2A* were generated from synthetic oligonucleotides. Four RGR–sgRNA modules were subsequently combined by Golden Gate assembly to produce the final pTVD-ECFP-RGR-STK11-CDKN2A-attB(Bxb1) donor containing four sgRNAs – two per target gene.

For MultiSite Gateway assembly, the entry/destination plasmid pTCV-attL4-attR1-attR2-attL3 was generated by Golden Gate assembly of synthetic oligonucleotides encoding the attL4 and attL3 Gateway sites together with the required BsaI cloning tandems into pTCV-BbsI. The complementary pTCVKanA-attR4-attR3 backbone/destination vector was generated in two steps. First, the attR4-ccdB-attR3 cassette was PCR-amplified from pDEST311 (Addgene plasmid 161896) and inserted into pTCV by Gibson assembly. A kanamycin-resistance cassette amplified from pCR2.1-TOPO was then introduced by Gibson assembly. PCR-amplified U6-sgRNA cassettes were inserted into the pTCV-attL4-attR1-attR2-attL3 vector by Golden Gate cloning to generate the intermediate entry/destination plasmid pTCV-attL4-sgRNA-CDKN2A-attR1-attR2-sgRNA-STK11-attL3, which was then used for MultiSite Gateway assembly with pENTR-mCherry-hygro and the appropriate pTCVKanA-attR4-attR3 Bxb1-attB destination backbone to generate pTVD-mCherry-hygro-sgRNA-CDKN2A-STK11-attB(Bxb1).

The Conditional Expression Cloning Cassette (CECC) was generated by sequential Gibson assembly of PCR-amplified fragments into pUC57. The cassette contained an EF1α promoter, a loxP-flanked transcriptional STOP module carrying the blasticidin-resistance and EYFP reporter genes, a multiple cloning site for insertion of the gene of interest, and a downstream T2A-RFP657 reporter. Gateway attB1 and attB2 sites flanked the complete cassette. A DNA fragment encoding HA-tagged KRASG12C was inserted into the CECC by KpnI/NotI restriction and ligation. The resulting construct was transferred into pDONR-zeo by Gateway BP recombination to generate the corresponding CECC-HA-KRAS^G12C^ entry clone. For MultiSite Gateway an intermediate entry/destination plasmid pTCV-attL4-attR1-attR2-attL3 containing WPRE-PolyA was generated by BsaI Golden Gate assembly using MTK6-015 (Addgene 123869_249819), MTK4-002 (123811_249751), 3 ds oligo adapters to provide compatible ends, and a ds oligo “plug” to block the left BsaI cloning tandem. These constructs were used for MultiSite Gateway recombination with the pTCVKanA-attR4-attR3 Bxb1-attB destination backbone to generate the pTVD-CE-KRAS^G12C^-attB(Bxb1) plasmid. All constructs were verified by diagnostic restriction digestion and Sanger sequencing as described above.

### PCR and digital PCR

Oligonucleotide sequences used for cloning, genotyping, and quantitative PCR are listed in Table S2. PCR fragments for Gibson and Golden Gate assembly were amplified using Q5 High-Fidelity DNA Polymerase (New England Biolabs, NEB), 0.5 µM each of forward and reverse primers, 0.2 mM each dNTP (Thermo Fisher Scientific), and 1–10 ng of plasmid DNA as template. For genotyping, genomic DNA (gDNA) was isolated from cultured cells using the DNeasy Blood & Tissue Kit (Qiagen) according to the manufacturer’s instructions. Genotyping PCR was performed using GoTaq G2 DNA Polymerase (Promega) with 100 ng of gDNA as template. PCR products used for cloning or Sanger sequencing were purified using the DNA Clean & Concentrator Kit (Zymo Research).

Transgene copy number was determined by duplex chip-based digital PCR (dPCR) using the QuantStudio 3D Digital PCR System (Applied Biosystems/Thermo Fisher Scientific). Genomic DNA was analyzed using a custom FAM-labeled Cas9-specific TaqMan assay and a VIC-labeled TaqMan Copy Number Reference Assay targeting the human RNase P (*RPPH1*) locus (both Applied Biosystems). Reactions were prepared using QuantStudio 3D Digital PCR reagents according to the manufacturer’s instructions, loaded onto digital PCR chips using the QuantStudio 3D Digital PCR Chip Loader, and amplified by thermal cycling. Chips were analyzed using the QuantStudio 3D Digital PCR Instrument and associated DigitalSuite/AnalysisSuite software. Cas9 copy number per diploid genome was calculated as triple the ratio of the Poisson-corrected Cas9 and RPPH1 concentrations, assuming three copies of *RPPH1* per HEK293 genome.

### Cell culture

HEK293 cells and their derivatives were maintained in Dulbecco’s modified Eagle medium (DMEM) supplemented with 10% fetal bovine serum (FBS; Anprotec), 100 U/mL penicillin, and 100 µg/mL streptomycin (Thermo Fisher Scientific) on tissue culture-treated plates (Sarstedt) at 37 °C in a humidified atmosphere containing 5% CO₂. Plasmid DNA was transfected using Lipofectamine 2000 (Thermo Fisher Scientific) and Opti-MEM reduced-serum medium (Thermo Fisher Scientific) according to the manufacturer’s instructions.

Human induced pluripotent stem cell (hiPSC) lines SCVI-15 and WTSIi004A were obtained from the Stanford Cardiovascular Institute Biobank and Sigma-Aldrich, respectively. Cells were maintained under feeder-free conditions on tissue culture plates (Sarstedt) coated with vitronectin (0.5 µg/cm²; Thermo Fisher Scientific) in Essential 8 (E8) medium (Thermo Fisher Scientific), which was replaced daily. For reverse transfection, hiPSCs were detached using StemMACS Passaging Solution XF (Miltenyi Biotec), collected in E8 medium supplemented with 10 µM ROCK inhibitor Y-27632 (MedChemExpress), and mixed with plasmid DNA–Lipofectamine Stem Transfection Reagent complexes (Thermo Fisher Scientific). Cells were tested for mycoplasma contamination using Mycoplasma PCR Detection Kit (Abcam).

TRAIT plasmids used for transfection are listed in the Table S1, other plasmids are pX459Blast, SGL40C.EFS.RFP657, pAAV-EF1a-Cre and pAAV-EF1a-Flpo (Addgene #118055, 69148, 55636, 55637).

For selection of genetically modified cells, HEK293 cells and their derivatives were cultured in DMEM supplemented with puromycin (1 µg/mL), G418 (500 µg/mL), hygromycin B (250 µg/mL), Zeocin (250 µg/mL), or blasticidin S (20 µg/mL), depending on the selectable marker. For hiPSCs, lower antibiotic concentrations were used: puromycin (0.25–1 µg/mL), G418 (100 µg/mL), hygromycin B (125 µg/mL), Zeocin (1.5 µg/mL), or blasticidin S (5-10 µg/mL).

### Puromycin-resistant colony-formation assay

Following plasmid transfection, 1 × 10⁵ cells were seeded into 10-cm tissue culture dishes and selected with puromycin (1 µg/mL) for 10 days. Surviving colonies were fixed with 70% ethanol, air-dried, and stained with 0.5% (w/v) crystal violet for 30 min at room temperature. Plates were subsequently rinsed with water to remove excess stain, air-dried, and scanned. Colonies were automatically counted using ImageJ (National Institutes of Health, USA).

### Immunofluorescence staining and fluorescence microscopy

For immunofluorescence detection of Cas9, cells were seeded into 8-well chamber slides (µ-Slide 8 Well, ibidi) and cultured in the presence or absence of doxycycline for 48 h. Cells were fixed with ice-cold methanol and blocked with 1% bovine serum albumin (BSA) in Tris-buffered saline containing 0.05% Tween-20 (TBS-T) for 1 h at room temperature. Primary and secondary antibodies were diluted in the same buffer. Cas9 was detected using mouse monoclonal anti-Cas9 antibody (clone 4G10; Diagenode, cat. no. C15200216; 1:100) incubated overnight at 4 °C, followed by DyLight 488-conjugated goat anti-mouse IgG (H+L) secondary antibody (Pierce/Thermo Fisher Scientific, cat. no. 35503; 1:200). Nuclei were counterstained with 300 nM 4′,6-diamidino-2-phenylindole (DAPI). Fluorescence images were acquired using a Leica DM5500 widefield fluorescence microscope (Leica Microsystems). For analysis of conditional fluorescent reporter expression in the Con/Fon assay, fluorescence was imaged directly in living cells cultured in tissue culture plates using an inverted Axio Observer.Z1 fluorescence microscope (Carl Zeiss Microscopy).

### Flow cytometry

Flow cytometry was used to quantify fluorescent reporter expression and expression of pluripotency markers, to assess the efficiency of transfection, and to detect recombinase-mediated cassette exchange. Cells were harvested and resuspended in MACS buffer. For pluripotency analysis primary antibodies FITC anti-human/mouse SSEA-3, PE anti-human TRA-1-81, PE/Cyanine7 anti-human TRA-1-60-R, and APC anti-human SSEA-4 (Biolegend cat. no 330321, 330418, 330708, 330620) and respective isotype controls (Biolegend cat. no 400805, 401609, 401628 and R&D Systems cat. no. IC007A) were used. After 30 min staining at 4C, cells were washed twice in MACS buffer and resuspended in PBS buffer containing 100 ng/mL DAPI to identify dead cells. Flow cytometric analysis was performed using a CytoFLEX LX flow cytometer (Beckman Coulter), and fluorescence-activated cell sorting (FACS) was performed using an Aurora CS Spectral Cell Sorter (Cytek Biosciences). GFP, ECFP, EYFP, mCherry, and RFP657 fluorescence was measured in living cells using appropriate excitation and detection channels. Reporter-negative control cells were used to establish fluorescence gates. Dead cells, identified by DAPI staining, and cellular debris were excluded from analysis. Where applicable, fluorescent reporter-positive populations were isolated by cell sorting. Flow cytometry data were analyzed using FlowJo software (version 10.10.1; BD Biosciences).

### Western blotting

Cells were washed with phosphate-buffered saline (PBS) and lysed in RIPA buffer supplemented with protease inhibitors (Roche). Lysates were incubated on ice for 15 min and clarified by centrifugation for 15 min at 4 °C. Protein concentrations were determined using the Bradford assay, and equal amounts of total protein were prepared for electrophoresis. Protein samples were mixed with 4× loading buffer and reducing agent, heated at 70 °C for 10 min, and separated on 12% Bis-Tris precast polyacrylamide gels using NuPAGE MES SDS Running Buffer (Invitrogen). A protein molecular-weight marker was included. Proteins were transferred onto Immun-Blot polyvinylidene difluoride (PVDF) membranes (0.2 µm pore size; Bio-Rad) by wet electrotransfer for 1 h at 4 °C using NuPAGE Transfer Buffer containing 20% methanol (Invitrogen). Membranes were blocked in Tris-buffered saline containing Tween-20 (TBS-T) and 5% nonfat dry milk. Primary antibodies were diluted in blocking buffer and incubated overnight at 4 °C. After three 10-min washes with TBS-T, membranes were incubated with the appropriate horseradish peroxidase (HRP)-conjugated secondary antibodies (1:2,000) for 1 h at room temperature. Following three additional 10-min washes with TBS-T, immunoreactive proteins were detected using SuperSignal West Dura chemiluminescent substrate (Thermo Fisher Scientific, cat. no. 34075). The following primary antibodies were used for WB: rabbit monoclonal anti-Cas9 (S. pyogenes; clone E7M1H; Cell Signaling Technology, cat. no. 19526; 1:500), mouse monoclonal anti-β-actin (clone AC-15; Abcam, cat. no. ab6276; 1:5,000), rabbit monoclonal anti-HA tag (clone C29F4; Cell Signaling Technology, cat. no. 3724; 1:500), and rabbit polyclonal anti-GFP (Abcam, cat. no. ab290; 1:500). Secondary antibodies were goat anti-rabbit IgG (H+L), poly-HRP40-conjugated (Fitzgerald, cat. no. 43R-IG096PHRP; 1:2,000), and goat anti-mouse IgG Fc, HRP-conjugated (Thermo Fisher Scientific, cat. no. A16084; 1:2,000).

### Reagent and data availability

TRAIT plasmids are intended for distribution through Addgene.

### Declaration of generative AI and Ai-assisted technologies in the writing and coding process

During the preparation of this manuscript, the authors used Large Language Models (ChatGPT by OpenAI, Claude by Anthropic) to assist with coding and script troubleshooting, as well as for improvement of grammar, language and readability of the manuscript. All AI assisted code and text-outputs were critically reviewed, verified and edited by the authors, who take full responsibility for the final content of the manuscript and the analyses presented.

## Acknowledgments

This work was supported by BMFTR grants 161L0279A and 03LW0520K. Authors thank Genomics Core Facility (Institute of Molecular Oncology, Dr. A.Nist and Prof.Dr. T.Stiewe), FACS core Facility of the Medical Faculty (Philipps University of Marburg) for technical support, Amara Kramer, Chloe Mikolajczyk, Ethan Hurt and Johanna Gutsmann for their contributions to molecular cloning, and Jana Schlochtermeier and Lilly Lamers for cell culture work, genome-engineering experiments, and molecular analyses. We gratefully acknowledge their contributions to the development and experimental validation of the TRAIT toolbox.

## Author contributions

JPH – conceptualization, experiments, data analysis, writing - editing; KA, PS – molecular biology and cell culture experiments, writing – editing; OT - funding acquisition, conceptualization, methodology, data curation, formal analysis, supervision, visualization, writing – original draft, editing.

## Abbreviations

4-OHT: 4-hydroxytamoxifen
CECC: Conditional Expression Cloning Cassette
CRISPR: clustered regularly interspaced short palindromic repeats
dPCR: digital polymerase chain reaction
DTS: DNA nuclear targeting sequence
ECFP: enhanced cyan fluorescent protein
EGFP: enhanced green fluorescent protein
EYFP: enhanced yellow fluorescent protein
FACS: fluorescence-activated cell sorting
FS: FrameSelect
GG: Golden Gate
GW: Gateway
HDR: homology-directed repair
HDV: hepatitis delta virus
HITI: homology-independent targeted integration
HH: hammerhead ribozyme
ICE: Inference of CRISPR Edits
iCG: inducible Cas9-GFP
iCGC: inducible Cas9-GFP-Cre
iPSC: induced pluripotent stem cell
LP: landing pad
MCS: multiple cloning site
MTK: Mammalian ToolKit
NHEJ: nonhomologous end joining
ORF: open reading frame
PCR: polymerase chain reaction
RGR: ribozyme-guide RNA-ribozyme
RMCE: recombinase-mediated cassette exchange
rtTA: reverse tetracycline-controlled transactivator
sgRNA: single-guide RNA
SSC: single-cell-derived clone
TRAIT: Toolbox for Rapid Assembly of Integration Templates
TRE: tetracycline-responsive element
WPRE: woodchuck hepatitis virus post-transcriptional regulatory element

