## Supplemental figures and tables for "TRAIT: A Toolbox for Rapid Assembly of Integration Templates for CRISPR/Cas- and Integrase-Mediated Genome Engineering"

### Supplementary materials

### Supplementary figure legends

**Supplementary Figure S1.** Generation and evaluation of TRAIT donor templates for HDR- and HITI-mediated targeted integration.

(A) Schematic representation of the experimental strategy for targeted integration into the human *hROSA26/THUMPDS3-AS1* locus. The GFP–puromycin-resistance cassette was incorporated into either a homology-directed repair (HDR) donor containing locus-specific homology arms (pTVH-hR26-GFP-puro) or a homology-independent targeted integration (HITI) donor containing FrameSelect recognition sequences and flanking regulatory elements (pTVF-ID-ULPF-GFP-puro). The corresponding sgRNA-expression constructs were used together with Cas9 to induce genomic cleavage and, in the HITI workflow, cleavage of the donor plasmid.

(B) Representative flow-cytometry histograms showing GFP, mCherry, and RFP657 expression in HEK293 cells following cotransfection with five plasmids (px459Blast for Cas9 expression, pTVF-ID-GFP-puro, two SGL40C.EFS.RFP657-sgRNA1/2 vectors targeting *hROSA26* and a pTVF-g0-mCherry plasmid expressing FrameSelector sgRNA for excision of the GFP-puro fragment). Fluorescence profiles are shown relative to nontransfected cells and the indicated individual reporter-containing constructs. The fractions of reporter-positive cells are indicated within the respective histograms.

(C) Representative images of puromycin-resistant colonies obtained with HDR- or HITI-compatible GFP–puromycin donor constructs targeting *hROSA26* or *AAVS1*, transfection was done as described in (A). For random integration control cells were transfected with pTVF-ID-GFP-puro vector only, without sgRNA and Cas9. Colonies were counted using ImageJ software, colony-detection plots are displayed below the corresponding plate images.

**Supplementary Figure S2.** Evaluation of doxycycline-inducible Cas9 expression and CreERT2 recombination activity in HEK293 cells.

(A) Representative flow-cytometry histograms showing EGFP expression following transient transfection of HEK293 cells with the inducible Cas9 constructs pTVF-ID-iCG and pTVF-ID-iCGC. Cells were cultured with or without doxycycline, as indicated. The right panels show overlays of fluorescence distributions from nontransfected cells, transfected cells without doxycycline, and doxycycline-treated transfected cells. The percentages indicate the EGFP-positive fractions within the analyzed populations.

(B) Schematic representation of the INTERSECT Con/Fon reporter, which requires the combined activity of Cre and Flpo recombinases for activation of mScarlet expression.

(C) Representative flow-cytometry histograms showing GFP and mScarlet expression in HEK293-iCGC cells (upper panels) and parental HEK293 cells (lower panels) following transfection with the Con/Fon reporter alone or together with a Flpo-expression plasmid. Where indicated, cells were treated with 1 µM 4-hydroxytamoxifen (4-OHT) for 24 h.

(D) Representative fluorescence microscopy images of parental HEK293 and HEK293-iCGC cells following transfection with the Con/Fon reporter and Flpo-expression plasmid. Reporter fluorescence illustrates recombinase-dependent activation in the HEK293-iCGC cells. Scale bars are shown in the images.

**Supplementary Figure S3.** PCR-based AAVS1 integration-junction analysis.

(A) Schematic representation of the targeting strategy and junction PCR. Primers for detection of the left and right junctions are shown in blue, primers for nested PCR used for amplification and sequencing of junction fragments are shown in red.

(B) PCR analysis of the *AAVS1* region targeted with the ECFP-zeo construct. DNA fragments were amplified with junction-specific PCRs, purified and used as template for nested PCRs shown in Figure

3, followed by Sanger sequencing shown in (C,D). No template (NTC) and gDNA from HEK293 and non-transfected HEK293-iCGC cells were used as controls.

(C,D) Representative Sanger sequencing chromatograms of the integration-specific PCR product, showing the donor–genome junction sequence at *AAVS1* locus. (C) DNA fragments from nested PCR were obtained using the left junction PCR fragment as the template (B) and sequenced with the locus-specific primer 9451 (top panel) and donor-specific primer 8254 (bottom panel). (D) DNA fragments from nested PCR were obtained using the right junction PCR fragment as the template (B) and sequenced using the locus-specific primer 15667 (top panel) and donor-specific primer 8254 (bottom panel). Mixed chromatograms after the junction point indicate presence of indels upon HITI.

**Supplementary Figure S4.** Design of TRAIT landing-pad vectors and validation of sgRNAs targeting *STK11* and *CDKN2A*.

(A) Schematic representation of the pTVF-ID-PaqCI vector for generation of HITI-compatible landing-pad integration templates. Two PaqCI Golden Gate cloning regions flank the Gateway attR1–ccdB–CmR–attR2 cassette and permit insertion of orthogonal Bxb1 attP attachment sites. FrameSelect recognition sequences, flanking insulator elements, and SV40 DNA nuclear targeting sequences (DTS) are also indicated.

(B) Schematic representation of pTVF-ID-BbsI-PaqCI, which combines PaqCI cloning sites for insertion of Bxb1 attP sequences with additional BbsI cloning sites for incorporation of locus-specific left and right homology arms (LHA and RHA). This configuration provides a donor-vector architecture suitable for preparing HDR- or HITI-compatible landing-pad templates.

(C) Validation of paired sgRNAs targeting exon 5 of *STK11* and exon 2 of *CDKN2A*. Agarose gel electrophoresis of PCR amplicons obtained from parental HEK293-iCGC cells and cells transfected with the respective Cas9/sgRNA constructs. For *STK11*, the predicted wild-type amplicon is 516 bp, whereas deletion of the approximately 90-bp region between the sgRNA sites generates a shorter product of approximately 430 bp. For *CDKN2A*, the expected wild-type product is 630 bp, and deletion of the approximately 40-bp region produces a shorter product of approximately 590 bp. An ApaLI restriction site located within the wild-type *CDKN2A* amplicon was used for additional analysis by restriction digestion. The positions of the sgRNA recognition sites and the deleted regions are shown beneath the gels.

(D) PCR analysis of the *STK11* and *CDKN2A* target regions in HEK293-iCGC cells following integration of the inducible RGR-sgRNA cassette, doxycycline treatment, and enrichment of reporter-positive cells by flow cytometry. Genomic DNA from parental HEK293-iCGC cells (lanes 1 and 4), doxycycline-treated, sorted HEK293-iCGC-igRNA cells (lanes 2 and 5), and no-template controls (lanes 3 and 6) was amplified using the indicated primer pairs. Expected amplicon sizes were 930 bp for *STK11* and 1,005 bp for *CDKN2A*. The highlighted bands indicate the amplified target regions used for subsequent analysis.

**Supplementary Figure S5.** Conditional KRAS$^{G12C}$ reporter expression and assembly of a Bxb1-compatible donor using MTK-derived regulatory elements and MultiSite Gateway recombination.

(A) Flow-cytometry analysis of EYFP and RFP657 expression in seven blasticidin-selected HEK293-iCGC clones carrying the conditional HA-KRAS$^{G12C}$ expression cassette. Cells were treated with 1 µM 4-hydroxytamoxifen (4-OHT) for 24 h before analysis. Fluorescence distributions of the individual clones are shown together with the parental HEK293-iCGC control. EYFP expression is associated with the unrecombined conditional cassette, whereas RFP657 expression reports activation of the downstream HA-KRAS$^{G12C}$–T2A–RFP657 expression module.

(B) Seven-fragment Golden Gate assembly of MTK-derived regulatory elements into the TRAIT MultiSite Gateway intermediate pTCV-attL4-attR1-attR2-attL3. The assembly comprises the vector backbone (fragment 1), three double-stranded adapter oligonucleotides (fragments 2, 4, and 6), an MTK-derived poly(A) element (fragment 3), an MTK-derived WPRE element (fragment 5), and a double-stranded oligonucleotide module (fragment 7). The adapters provide compatible assembly junctions and introduce PaqCI recognition sites flanking the WPRE–poly(A) module. Fragment 7 replaces the original

BsaI cloning tandem with BbsI cloning sites, preserving the possibility of subsequent Golden Gate modification using a different Type IIS restriction enzyme.

(C) MultiSite Gateway assembly of a Bxb1-compatible conditional KRAS$^{G12C}$ donor. Three independently prepared components were combined: the existing pENTR-CE-HA-KRASG12C-RFP657 expression cassette, the pTCV-attL4-attR1-attR2-attL3-WPRE-pA intermediate generated in panel B, and the pTCVKanA-attR4-attR3-attB(Bxb1) destination backbone carrying orthogonal Bxb1 attachment sites. The resulting construct, pTVD-CE-HA-KRASG12C-attB(Bxb1), contains the conditional-expression cassette, WPRE–poly(A) regulatory elements, and Bxb1-compatible integration sites. Correct assembly was verified by diagnostic restriction digestion and Sanger sequencing of selected fragments and assembly junctions.

**Supplementary Figure S6.** Molecular screening and characterization of human iPSC populations engineered with TRAIT-derived inducible Cas9 and landing-pad constructs.

(A) PCR analysis of inducible Cas9 (iCG) donor sequences and integration junctions in SCVI-15 (upper panels) and WTSIi004A (lower panels) human iPSC populations following HITI-mediated targeting. Internal PCR assays (left panels) detect the neomycin-resistance (NeoR) and Cas9 sequences, whereas locus-specific PCR (right panels) assays examine integration at the *SHS231* locus. The SCVI-15 analysis includes an assay for antisense-orientation integration, while the WTSI analysis includes assays for both left and right SHS231 integration junctions. Parental iPSCs, engineered populations, no-template controls (NTC), and a HEK293-iCGC positive control are included as indicated. Primer combinations and expected amplicon sizes are shown adjacent to the gels.

(B) Representative flow-cytometry histograms showing doxycycline-treated SCVI-15-iCG and WTSI-iCG cells from (A) with confirmed targeted integration in *SHS231* locus. No GFP expression is detected after induction. Nontransfected cells and cells maintained without doxycycline are shown as controls.

(C) Representative immunofluorescence images showing doxycycline-inducible Cas9 expression in selected WTSI-iCG clones with confirmed targeted integration in *hROSA26* locus. Cells were treated with doxycycline (1 µg/mL) for 48 h. Cas9 immunostaining is shown in green, and nuclei are counterstained with DAPI (blue). Scale bar, 125 µm.

(D) PCR amplification of the TP53 exon 3 and exon 5 target regions in parental WTSI-iCG cells, and the Nutlin-3A-selected WTSI-iCG TP53-edited population. No-template controls are included.

(E) Representative Sanger sequencing chromatograms of the TP53 exon 3 and exon 5 target regions from the Nutlin-3A-selected WTSI-iCG population and corresponding control samples. The chromatograms illustrate sequence alterations in the selected population and provide the input for ICE-based editing analysis presented in Figure 7F.

**Supplementary Figure S7.** Molecular screening and characterization of WTSI-iCG cells with stable integration of landing-pad constructs.

(A) Flow-cytometric analysis of fluorescent reporter expression in SCVI-15 iPSCs following introduction of TRAIT-derived landing-pad constructs. Fluorescence distributions before and after antibiotic selection are shown for the EYFP–blasticidin, mCherry–hygromycin, and ECFP–zeocin constructs.

(B) PCR analysis of antibiotic-selected, FACS-sorted SCVI-15 cells transfected with indicated landing pad constructs. PCR using internal primers specific for landing pads (left panel – left flank, right panel – right flank) was used to confirm presence of the constructs in pools of selected sorted cells (S1, S2 - two independently generated cell pools per landing pad).

(C) Schematic representation of the landing pad and PCR strategy used in (B).

**Supplementary Table 1 TRAIT plasmids**

**Supplementary_table_1.xlsx**

**Supplementary Table 2 Oligonucleotides used for cloning and PCR**

**Supplementary_table_2.xlsx**

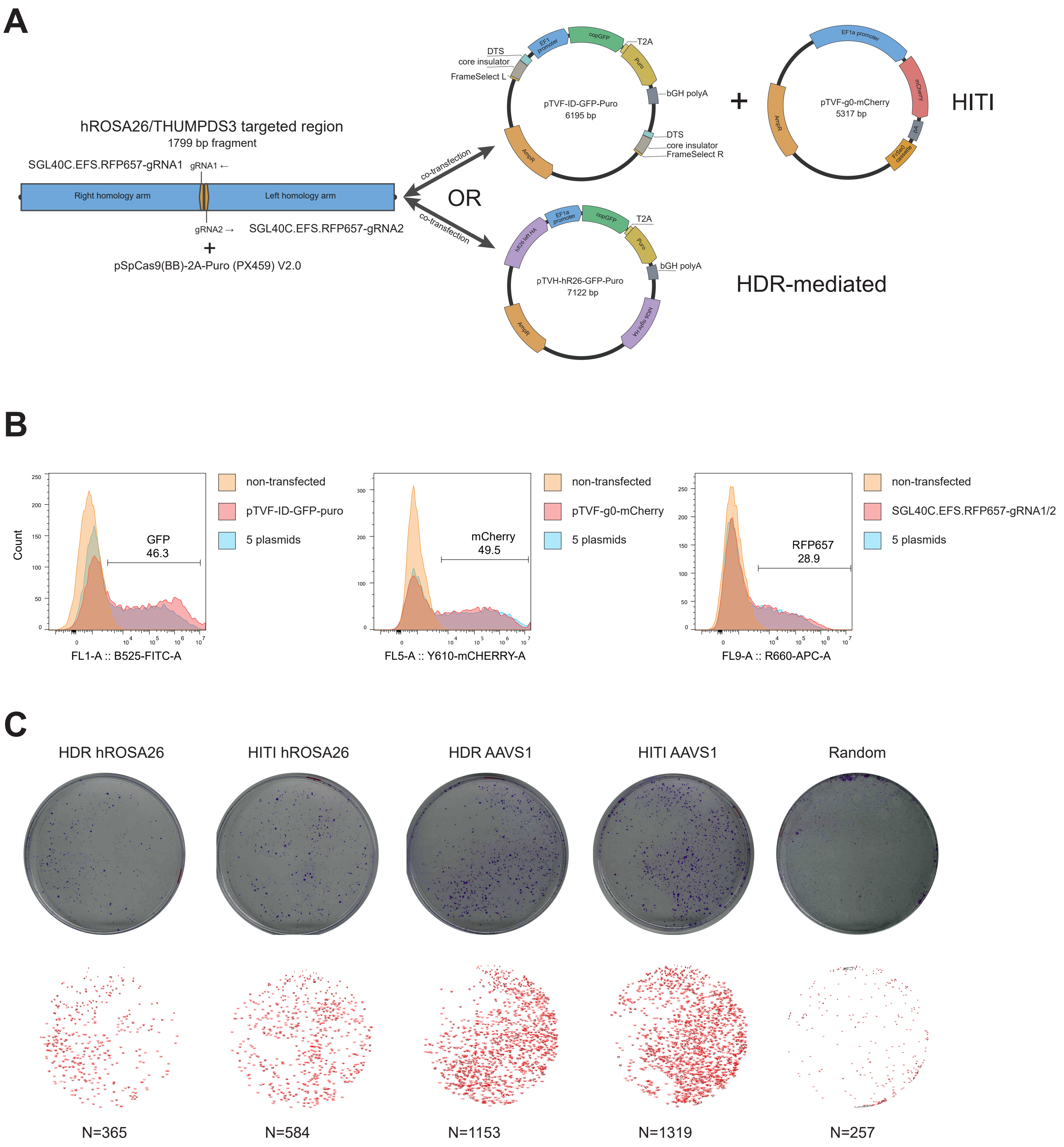


Supplementary Figure 1

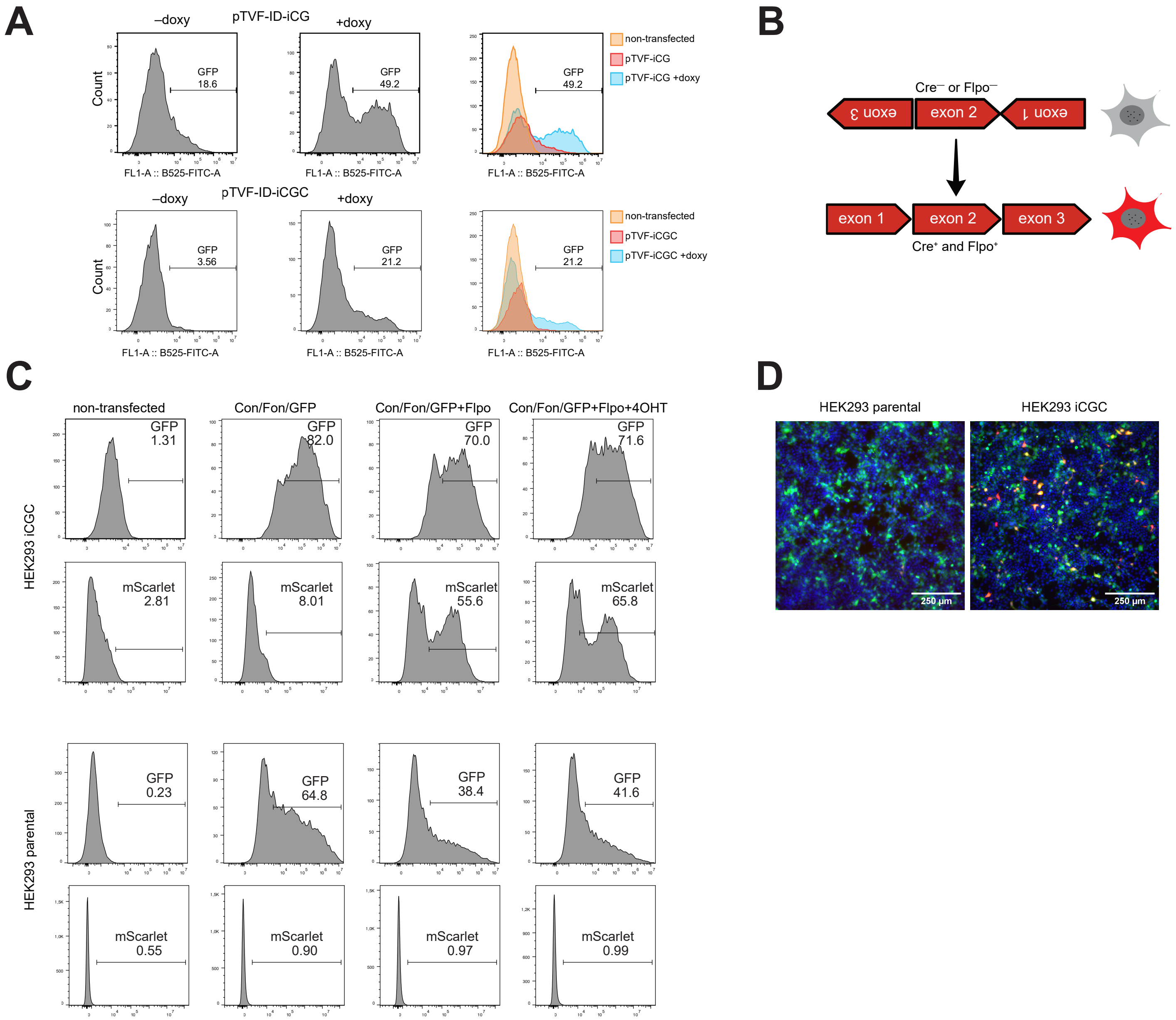


Supplementary Figure 2

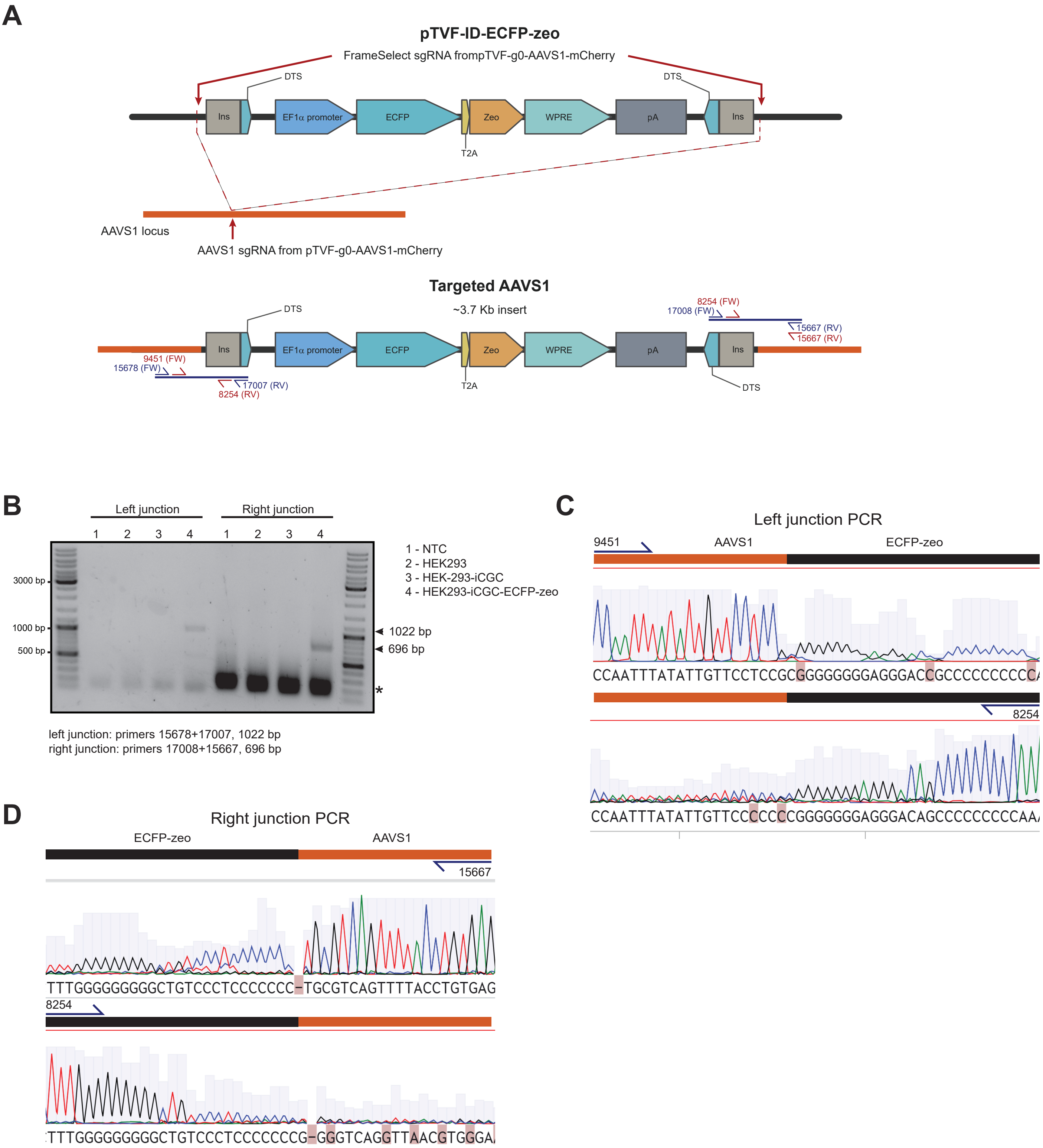
A
pTVF-ID-ECFP-zeo
FrameSelect sgRNA frompTVF-g0-AAVS1-mCherry
DTS
DTS
Ins
EF1α promoter
ECFP
Zeo
WPRE
pA
Ins
T2A
AAVS1 locus
AAVS1 sgRNA from pTVF-g0-AAVS1-mCherry
Targeted AAVS1
~3.7 Kb insert
8254 (FW)
17008 (FW)
15667 (RV)
15667 (RV)
9451 (FW)
15678 (FW)
17007 (RV)
8254 (RV)
DTS
DTS
B
Left junction
Right junction
1 2 3 4
1 2 3 4
3000 bp
1000 bp
500 bp
1 - NTC
2 - HEK293
3 - HEK-293-iCGC
4 - HEK293-iCGC-ECFP-zeo
1022 bp
696 bp
*
left junction: primers 15678+17007, 1022 bp
right junction: primers 17008+15667, 696 bp
C
Left junction PCR
9451
AAVS1
ECFP-zeo
CCAATTTATATTGTTCCTCCGCGGGGGGGGAGGGACCGCCCCCCCCCCA
8254
CCAATTTATATTGTTCCCCCCCGGGGGGGAGGGACAGCCCCCCCCCAA
D
Right junction PCR
ECFP-zeo
AAVS1
15667
TTTGGGGGGGGGCTGTCCCTCCCCCCCC-TGCGTCAGTTTTACCTGTGAG
8254
TTTGGGGGGGGGCTGTCCCTCCCCCCCCG-GGGTCAGGTTAACGTGGGA

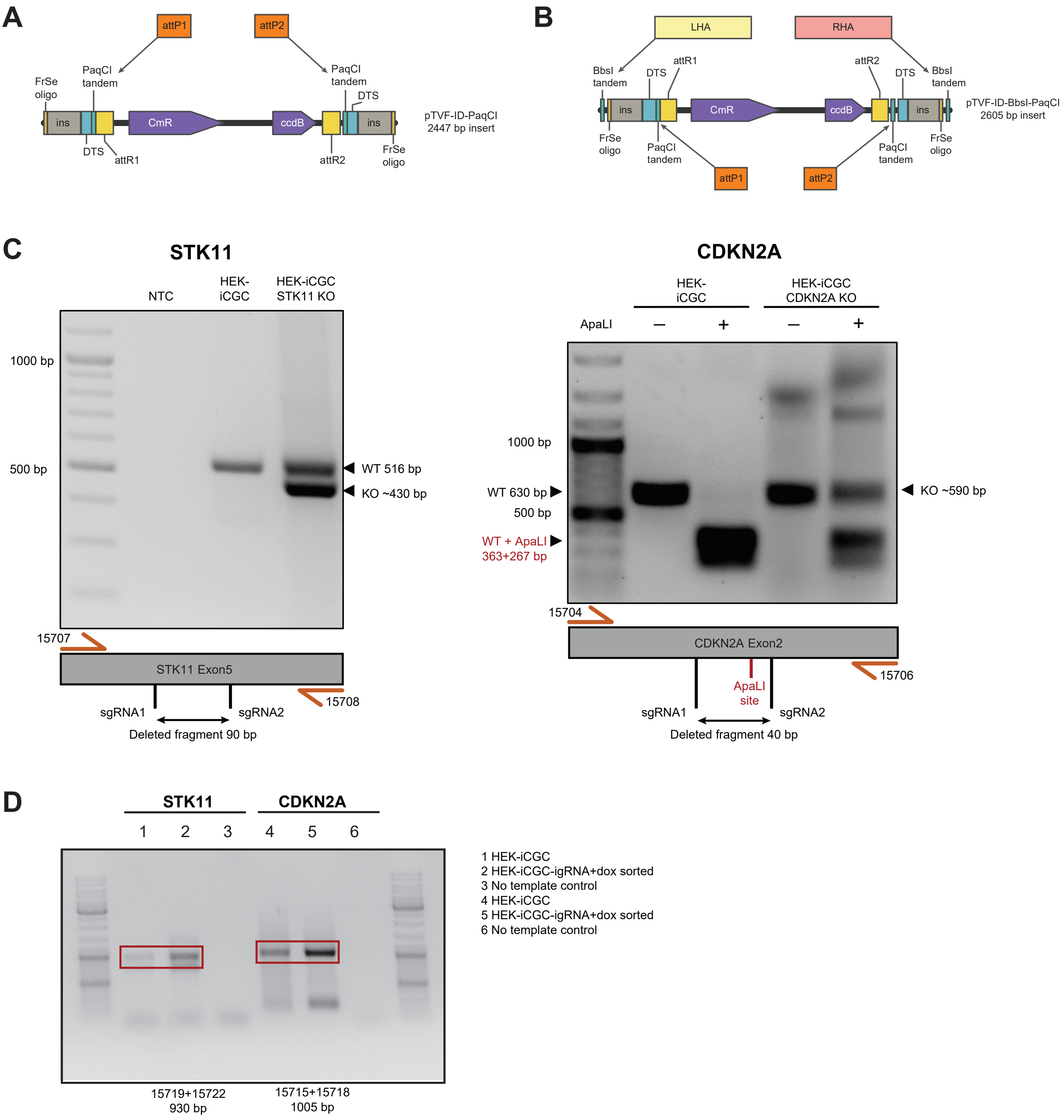


Supplementary Figure 4

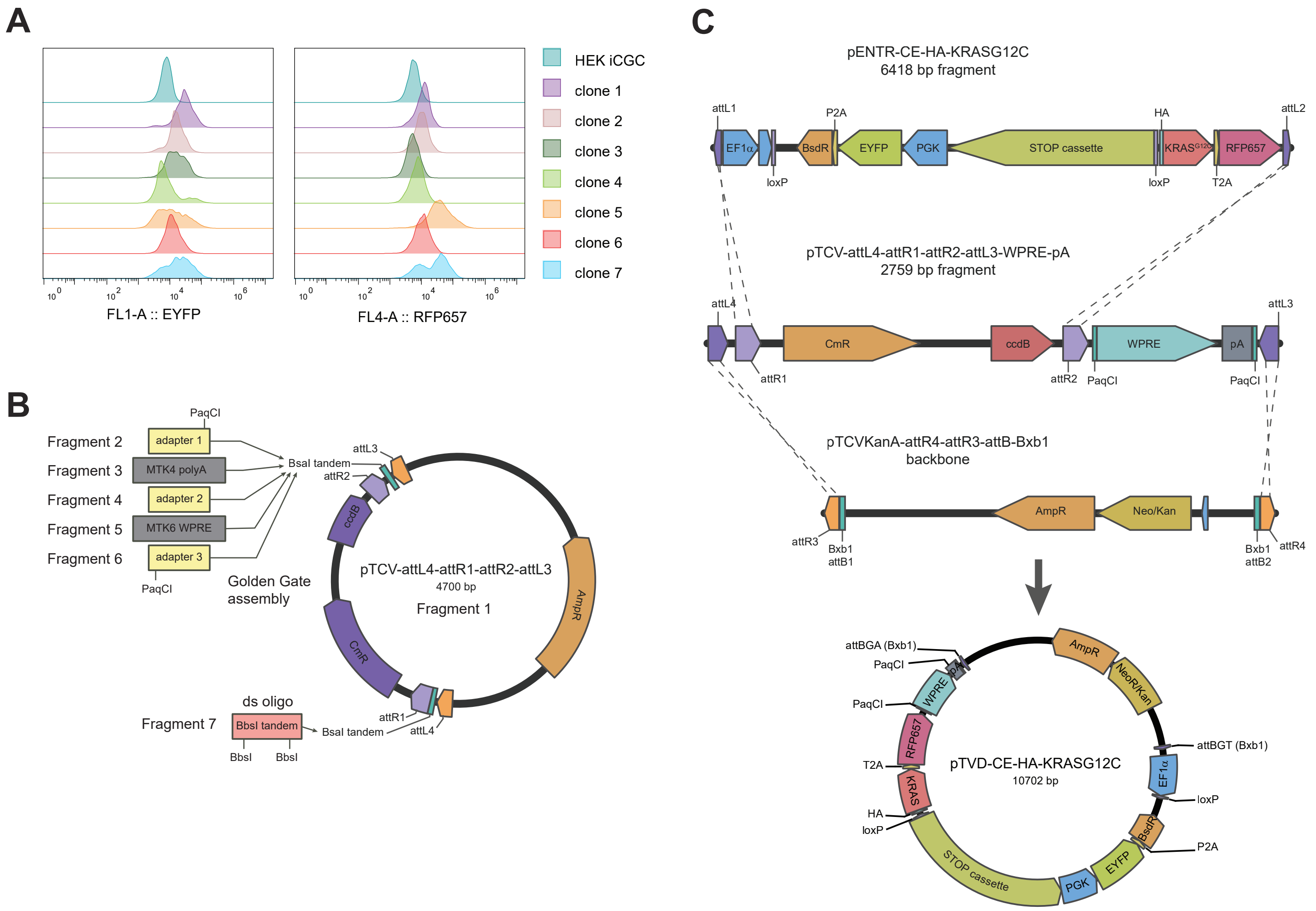


Supplementary Figure 5

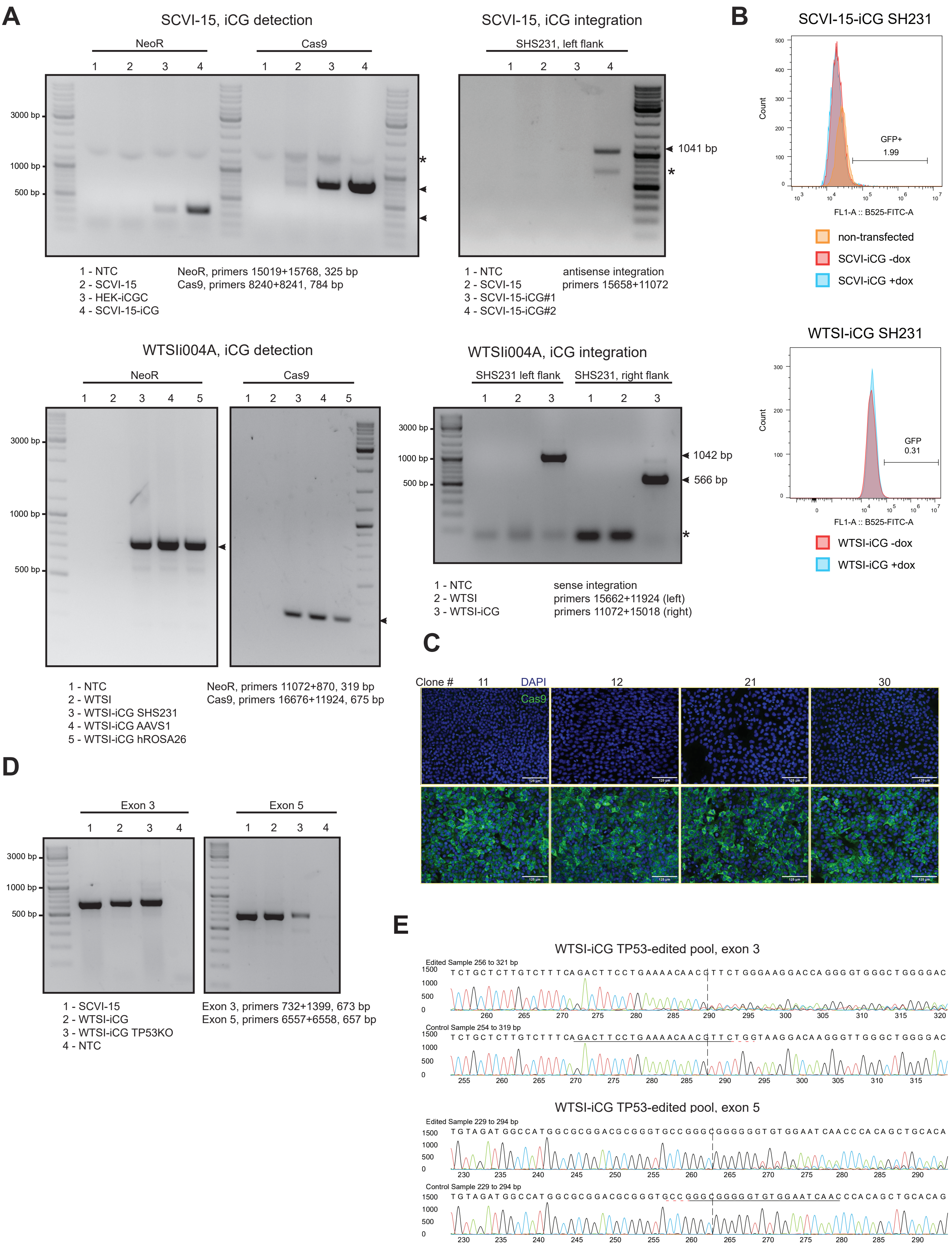


Supplementary Figure 6

A

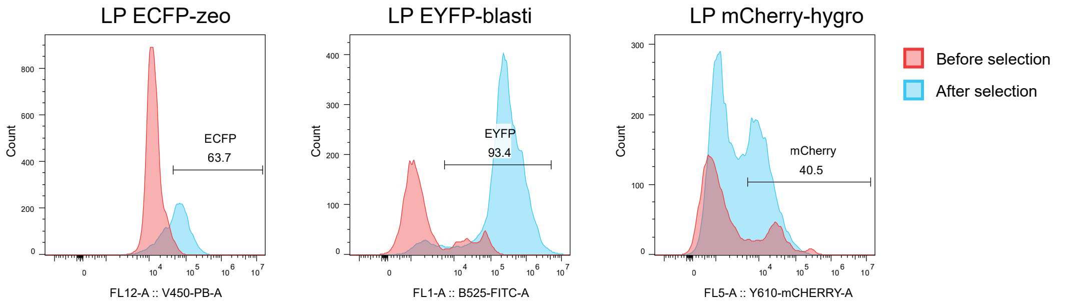
LP ECFP-zeo
LP EYFP-blasti
LP mCherry-hygro
Before selection
After selection
Count
ECFP
63.7
EYFP
93.4
mCherry
40.5
FL12-A :: V450-PB-A
FL1-A :: B525-FITC-A
FL5-A :: Y610-mCHERRY-A

B

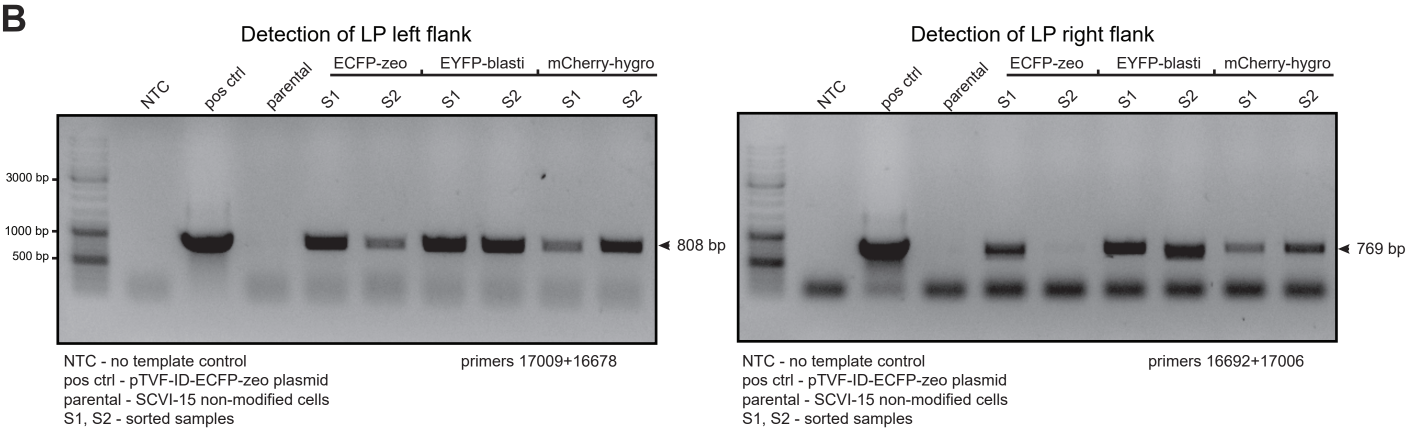
Detection of LP left flank
NTC
pos ctrl
parental
ECFP-zeo
EYFP-blasti
mCherry-hygro
S1
S2
3000 bp
1000 bp
500 bp
808 bp
NTC - no template control
pos ctrl - pTVF-ID-ECFP-zeo plasmid
parental - SCVI-15 non-modified cells
S1, S2 - sorted samples
primers 17009+16678
Detection of LP right flank
769 bp
primers 16692+17006

C

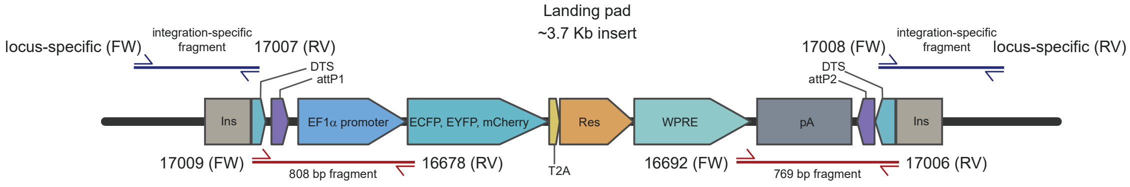
Landing pad
~3.7 Kb insert
locus-specific (FW)
integration-specific fragment
17007 (RV)
DTS
attP1
17008 (FW)
attP2
locus-specific (RV)
Ins
EF1α promoter
ECFP, EYFP, mCherry
Res
WPRE
pA
17009 (FW)
808 bp fragment
16678 (RV)
T2A
16692 (FW)
769 bp fragment
17006 (RV)

**Supplementary Table 1. TRAIT plasmids**

| Plasmid ID | Plasmid name | Design / application |
|---|---|---|
| G284 | pTCV-BsaI-free | Core Template Cloning Vector (pTCV) backbones |
| G764 | pTCV-BbsI-only | Core Template Cloning Vector (pTCV) backbones |
| G765 | pTCV-BsaI-only | Core Template Cloning Vector (pTCV) backbones |
| H097 | pTCV-PaqCI | Core Template Cloning Vector (pTCV) backbones |
| G778 | pTVF | FrameSelect, insulator and landing-pad-ready donor backbones |
| G863 | pTVFI | FrameSelect, insulator and landing-pad-ready donor backbones |
| G866 | pTVFD | FrameSelect, insulator and landing-pad-ready donor backbones |
| G867 | pTVF-ID | FrameSelect, insulator and landing-pad-ready donor backbones |
| H213 | pTVF-ID-BbsI-PaqCI | FrameSelect, insulator and landing-pad-ready donor backbones |
| G413 | pTCV-hROSA26-HITI | Locus-targeted HDR and HITI donors |
| G789 | pTCV-SHS231-HITI | Locus-targeted HDR and HITI donors |
| H464 | pTVH-AAVS1 | Locus-targeted HDR and HITI donors |
| H465 | pTVH-hROSA26 | Locus-targeted HDR and HITI donors |
| H466 | pTVH-AAVS1-GFP-puro | Locus-targeted HDR and HITI donors |
| H467 | pTVH-ROSA26-GFP-puro | Locus-targeted HDR and HITI donors |
| G771 | pTVF-g0-BsaI | FrameSelect and genomic-targeting sgRNA vectors |
| G772 | pTVF-g+1-BsaI | FrameSelect and genomic-targeting sgRNA vectors |
| G773 | pTVF-g+2-BsaI | FrameSelect and genomic-targeting sgRNA vectors |
| G774 | pTVF-g-BbsI | FrameSelect and genomic-targeting sgRNA vectors |
| G775 | pTVF-g+1-BbsI | FrameSelect and genomic-targeting sgRNA vectors |
| G776 | pTVF-g+2-BbsI | FrameSelect and genomic-targeting sgRNA vectors |
| G782 | pTVF-go0-BbsI | FrameSelect and genomic-targeting sgRNA vectors |
| G783 | pTVF-go+1-BbsI | FrameSelect and genomic-targeting sgRNA vectors |
| G784 | pTVF-go+2-BbsI | FrameSelect and genomic-targeting sgRNA vectors |
| G785 | pTVF-go0-BsaI | FrameSelect and genomic-targeting sgRNA vectors |
| G786 | pTVF-go+1-BsaI | FrameSelect and genomic-targeting sgRNA vectors |
| G787 | pTVF-go+2-BsaI | FrameSelect and genomic-targeting sgRNA vectors |
| G791 | pTVF-go0-BsaI-mCherry | FrameSelect and genomic-targeting sgRNA vectors |
| G792 | pTVF-g0-BsaI-mCherry | FrameSelect and genomic-targeting sgRNA vectors |
| G793 | pTVF-g+1-BsaI-mCherry | FrameSelect and genomic-targeting sgRNA vectors |
| G794 | pTVF-g+2-BsaI-mCherry | FrameSelect and genomic-targeting sgRNA vectors |
| G795 | pTVF-go+2-BsaI-mCherry | FrameSelect and genomic-targeting sgRNA vectors |
| G796 | pTVF-go+1-BsaI-mCherry | FrameSelect and genomic-targeting sgRNA vectors |
| G839 | pTVF-go0-hR26-1-mCherry | FrameSelect and genomic-targeting sgRNA vectors |
| G840 | pTVF-go0-hR26-2-mCherry | FrameSelect and genomic-targeting sgRNA vectors |
| G841 | pTVF-go0-AAVS1-1-mCherry | FrameSelect and genomic-targeting sgRNA vectors |
| G842 | pTVF-go0-AAVS1-2-mCherry | FrameSelect and genomic-targeting sgRNA vectors |
| G843 | pTVF-go0-SHS231-1-mCherry | FrameSelect and genomic-targeting sgRNA vectors |
| G844 | pTVF-go0-SHS231-2-mCherry | FrameSelect and genomic-targeting sgRNA vectors |
| G761 | pUC57-iCGCv1 | Inducible Cas9 integration and Gateway entry constructs |
| G762 | pENTR-iCGCv1 | Inducible Cas9 integration and Gateway entry constructs |
| G763 | pTCV-iCGCv1-SHS231-HITI | Inducible Cas9 integration and Gateway entry constructs |
| G790 | pTVF-iCGCv1 | Inducible Cas9 integration and Gateway entry constructs |
| G814 | pUC57-iCG | Inducible Cas9 integration and Gateway entry constructs |
| G819 | pENTR-iCG | Inducible Cas9 integration and Gateway entry constructs |
| G825 | pTVF-iCG | Inducible Cas9 integration and Gateway entry constructs |
| G864 | pTVFI-iCG | Inducible Cas9 integration and Gateway entry constructs |
| G983 | pTVF-ID-gRNA-LP | Bxb1 landing-pad and inducible gRNA-LP constructs |
| G994 | pTVF-ID-gRNA-LP-attP(Bxb1) | Bxb1 landing-pad and inducible gRNA-LP constructs |
| I192 | pTVF-ID-BbsI-attP(Bxb1) | Bxb1 landing-pad and inducible gRNA-LP constructs |
| I196 | pTVF-ID-LP-GFP-puro-attP(Bxb1) | Bxb1 landing-pad and inducible gRNA-LP constructs |
| G969 | pENTR-ECFP-zeo | RGR editing donors and Gateway reporter/selection entries |
| G979 | pENTR-GFP-puro | RGR editing donors and Gateway reporter/selection entries |
| H020 | pTVD-ECFP-RGR-STK11-CDKN2A-attB(Bxb1) | RGR editing donors and Gateway reporter/selection entries |
| H116 | pENTR-mCherry-Hygro | MultiSite Gateway and Bxb1 destination/intermediate vectors |
| I193 | pTCVKanA-attR4-attR3-attB(Bxb1) | MultiSite Gateway and Bxb1 destination/intermediate vectors |
| I195 | pTCV-attL4-attR1-attR2-attL3-WPRE-pA | MultiSite Gateway and Bxb1 destination/intermediate vectors |
| G322 | pCECC | Conditional expression and oncogenic payloads |
| G417 | pCE-HA-KRASG12C | Conditional expression and oncogenic payloads |
| G476 | pCE-HA-KRASG12D | Conditional expression and oncogenic payloads |
| G477 | pENTR-CE-HA-KRASG12C | Conditional expression and oncogenic payloads |
| G596 | pCE-EGFRL858R | Conditional expression and oncogenic payloads |
| G688 | pENTR-CE-HA-KRASG12D | Conditional expression and oncogenic payloads |
| G695 | pENTR-CE-EGFRL858R | Conditional expression and oncogenic payloads |
| G698 | pCE-PIK3CAE545K | Conditional expression and oncogenic payloads |
| G752 | pENTR-CE-PIK3CAE545K | Conditional expression and oncogenic payloads |
| H113 | pTVF-ID-CE-KRASG12D | Conditional expression and oncogenic payloads |
| H114 | pTVF-ID-CE-KRASG12C | Conditional expression and oncogenic payloads |
| I198 | pTVD-CE-HA-KRASG12C-attB(Bxb1) | Conditional expression and oncogenic payloads |
| G689 | pTVH-AAVS1-HA-KRASG12C | Oncogene HDR donor collection |
| G690 | pTVH-hROSA26-HA-KRASG12C | Oncogene HDR donor collection |
| G691 | pTVH-AAVS1-HA-KRASG12D | Oncogene HDR donor collection |
| G692 | pTVH-hROSA26-HA-KRASG12D | Oncogene HDR donor collection |
| G696 | pTVH-AAVS1-EGFRL858R | Oncogene HDR donor collection |
| G697 | pTVH-hROSA26-EGFRL858R | Oncogene HDR donor collection |
| G753 | pTVH-AAVS1-PIK3CAE545K | Oncogene HDR donor collection |
| G754 | pTVH-hROSA26-PIK3CAE545K | Oncogene HDR donor collection |
| G788 | pENTR-mCherry-BsaI-free | Reusable reporter and accessory entry modules |
| G822 | pTVF-Con/Fon-oScarlet-puro-GFP | Reusable reporter and accessory entry modules |
| H353 | pENTR-RFP657-Puro | Reusable reporter and accessory entry modules |
| H354 | pENTR-RFP657-Hygro | Reusable reporter and accessory entry modules |
| H947 | pTCV-attL4-sgRNA-CDKN2A-attR1-attR2-sgRNA-STK11-attL3 | Intermediate donor/acceptor plasmid for MultiSite Gateway |

**Supplementary Table 2. Oligonucleotides used for cloning and PCR**

| NN | Oligonucleotide sequence | Oligo name | Oligo ID | Purpose | Vector | Extension | Type IIS sites | Left overhang, 5'->3' | Right overhang, 5'->3' |
|---|---|---|---|---|---|---|---|---|---|
| 1 | CCTggtctcaaaccGATCACGCGAGGAGGAAAGGAGGGAGG | BsaI-ROSA26_RV | 15376 | PCR primer GG cloning | pTVH-hROSA26 | | BsaI | | aacc |
| 2 | GGGgaagaccccaccCGCTTCTCGATTATGGGCGGGATTCTTTTGC | BbsI-ROSA26_FW | 15377 | PCR primer GG cloning | pTVH-hROSA26 | | BbsI | cacc | |
| 3 | CCCCGATAACGGAcACCGGCACACTGGCC | Mut-ccdB-V73_FW | 15413 | SDM ccdB primer | pTCV | | | | |
| 4 | GGCCAGTGTGCCGGTgTCCGTTATCGGGG | Mut-ccdB-V73_RV | 15414 | SDM ccdB primer | pTCV | | | | |
| 5 | ccagtgaattcgagctcggtacACAAGTTTGTACAAAAAAGCAGGCTAAGGATCTGCGATCGCTCC | GibAsI_F1_FW | 15419 | PCR primer Gibson Assembly | pUC57-CECC | | | | |
| 6 | cgaaaggcccggagatgagg | GibAsI_F4_FW | 15425 | PCR primer Gibson Assembly | pUC57-CECC | | | | |
| 7 | gcccgggatccgatatctagACCACTTTGTACAAGAAAGCTGGGTttaattaagcttgtgccccagtttgc | GibAsII_F6_RV_new | 15579 | PCR primer Gibson Assembly | pUC57-CECC | | | | |
| 8 | GGAGGTACCccggtgagaccgagagagggtctcaggttCAAGTTTGTACAAAAAAGC | KpnI-ccdB-2_FW | 15592 | PCR primer cloning | pTCV | KpnI | BsaI | ccgg | aacc |
| 9 | CCTTCTAGAaatgcaggtcttctcgaagacccggtgtGATCTGTCGAATCGATCACC | XbaI-ccdB-2_RV | 15593 | PCR primer cloning | pTCV | XbaI | BbsI | cacc | atgc |
| 10 | GGAggtctcTccggGTCCTGCTTTCTCTGACCTGC | BsaI-AAVS1_FW | 15664 | PCR primer GG cloning | pTVH-AAVS1 | | BsaI | ccgg | |
| 11 | CCTggtctcaaaccGCCCCACTGTGGGGTGGAGG | BsaI-AAVS1_RV | 15665 | PCR primer GG cloning | pTVH-AAVS1 | | BsaI | | aacc |
| 12 | GGGgaagaccccaccGGGACAGGATTGGTGACAGAAAAG | BbsI-AAVS1_FW | 15666 | PCR primer GG cloning | pTVH-AAVS1 | | BbsI | cacc | |
| 13 | GGGgaagactgatgcGAGCAGAGCCAGGAACCCCTG | BbsI-AAVS1_RV | 15667 | PCR primer GG cloning | pTVH-AAVS1 | | BbsI | | atgc |
| 14 | GGAggtctcTccggGACACCTGATTGGCTGTCGAAGC | BsaI-hR26_FW | 15727 | PCR primer GG cloning | pTVH-hROSA26 | | BsaI | ccgg | |
| 15 | GGGgaagactgatgcGGTATAAAACTCAGGTGATCATGC | BbsI-hR26_RV | 15728 | PCR primer GG cloning | pTVH-hROSA26 | | BbsI | | atgc |
| 16 | ctagatatcggatcccgggc | pUC57-XbaI_FW | 15819 | PCR primer Gibson Assembly | pUC57-mCherry | | | | |
| 17 | GTACcgagctcgaattcactgg | pUC57-KpnI_RV | 15820 | PCR primer Gibson Assembly | pUC57-mCherry | | | | |
| 18 | caatgataccgcgagaTccacgctcacc | Mut-Amp-G242_FW | 15823 | SDM Amp primer | pTCV | | | | |
| 19 | caatgataccgcgagatccacgctcacc | Mut-Amp-G242_FW | 15823 | PCR primer Gibson Assembly | pTVF-ID-gRNA-LP | | | | |
| 20 | ggtgagcgtggAtctcgcggtatcattg | Mut-Amp-G242_RV | 15824 | SDM Amp primer | pTCV | | | | |
| 21 | ggtgagcgtggatctcgcggtatcattg | Mut-Amp-G242_RV | 15824 | PCR primer Gibson Assembly | pTVF-ID-gRNA-LP | | | | |
| 22 | ggtgagcgtggatctcgcggtatcattg | Mut-Amp-G242_RV | 15824 | PCR primer Gibson Assembly | pTCVKan-attR4-attR3-PaqCI | | | | |
| 23 | ctcctcttcctcatctccgggcctttcgGCCACCATGGTGAGCAAGGG | GibAsInew_F1_RV | 16133 | PCR primer Gibson Assembly | pUC57-CECC | | | | |
| 24 | ccttcacaaagatccctcgaccagacatactcgaccagacatataagatac | GibAsC_F2_RV | 16134 | PCR primer Gibson Assembly | pUC57-CECC | | | | |
| 25 | tatgtctggtcgagggatctttgtg | GibAsC_F3_FW | 16135 | PCR primer Gibson Assembly | pUC57-CECC | | | | |
| 26 | cgacggccagtgaattcgagctcggtacacaagtttgtacaaaaaagcaggctgaggatcctttaccactccc | GAs-iCGC-F1_FW | 16676 | PCR primer Gibson Assembly | iCGC | attB1 | | | |
| 27 | gattacaaagacgatgacgataagatggcc | GAs-iCGC-F2_FW | 16677 | PCR primer Gibson Assembly | iCGC | | | | |
| 28 | tcctcgcccttgctcaccat | GAs-iCGC-F2_RV | 16678 | PCR primer Gibson Assembly | iCGC | | | | |
| 29 | ccccgaattcatggtgagcaagggcgagga | GAs-iCGC-F3_FW | 16679 | PCR primer Gibson Assembly | iCGC | | | | |
| 30 | tcacttgtacagctcgtccatg | GAs-iCGC-F3_RV | 16680 | PCR primer Gibson Assembly | iCGC | | | | |
| 31 | cactctcggcatggacgagctgtacaagtgaggcagtctggagtaacccacccaagatctgg | GAs-iCGC-F4_FW | 16681 | PCR primer Gibson Assembly | iCGC | | | | |
| 32 | cCtctagtGcgGgtCgaCTTAtgtg | GAs-iCGC-F4_RV | 16682 | PCR primer Gibson Assembly | iCGC | | | | |
| 33 | cctgccacaTAAGtcGacCcgCactagaGg | GAs-iCGC-F5_FW | 16683 | PCR primer Gibson Assembly | iCGC | | | | |
| 34 | cagtcgacgggcccgggatccgatatctagACCACTTTGTACAAGAAAGCTGGGTtaagatacattgatgagtttggacaaaccac | GAs-iCGC-F5_RV | 16684 | PCR primer Gibson Assembly | iCGC | attB2 | | | |
| 35 | ggccatcttatcgtcatcgtctttgtaatcccatggtggcccaggcgatctgacggttcactaaacgagc | GAs-iCGC-F1_RV | 16685 | PCR primer Gibson Assembly | iCGC | | | | |
| 36 | ccggacgtcttcgagaagaccg | BsaI-to-BbsI_S | 16730 | BsaI to BbsI GG oligo | pTCV-BbsI | | BbsI | ccgg | aacc |
| 37 | aacccggtcttctcgaagacgt | BsaI-to-BbsI_AS | 16731 | BsaI to BbsI GG oligo | pTCV-BbsI | | BbsI | ccgg | aacc |
| 38 | cacccgagaccgagagagggtctcc | BbsI-to-BsaI_S | 16732 | BbsI to BsaI GG oligo | pTCV-BsaI | | BsaI | cacc | atgc |
| 39 | atgcggagaccctctctcggtctcg | BbsI-to-BsaI_AS | 16733 | BbsI to BsaI GG oligo | pTCV-BsaI | | BsaI | cacc | atgc |
| 40 | GAGTTTTTGTTCAGGGCCGGGATTCTCTTCCAC | GAs-iCGDC-F3_RV | 16734 | PCR primer Gibson Assembly | iCGC | | | | |
| 41 | GAAGAGAATCCCGGCCCTGAACAAAAACTCATCTCAGAAGAGGATCTG | GAs-iCGDC-F4_FW | 16735 | PCR primer Gibson Assembly | iCGDC | | | | |
| 42 | gtaaattggaccgctccagaatctcaaagcaatagc | GAs-iCGDC-F4_RV | 16736 | PCR primer Gibson Assembly | iCGDC | | | | |
| 43 | gctttgagattctggagcggtccaatttactgaccgtacaccaaaatttg | GAs-iCGDC-F5_FW | 16737 | PCR primer Gibson Assembly | iCGDC | | | | |
| 44 | CACCgccagtacccaaaaagcggg | FrSe0-gRNA_S | 16738 | gRNA GG oligo | pX459 | | | cacc | |
| 45 | AAACcccgctttttgggtactggc | FrSe0-gRNA_AS | 16739 | gRNA GG oligo | pX459 | | | | aaac |
| 46 | CACCggccagtacccaaaaagcgg | FrSe1-gRNA_S | 16740 | gRNA GG oligo | pX459 | | | cacc | |
| 47 | AAACccgctttttgggtactggcc | FrSe1-gRNA_AS | 16741 | gRNA GG oligo | pX459 | | | | aaac |
| 48 | CACCgggccagtacccaaaaagcg | FrSe2-gRNA_S | 16742 | gRNA GG oligo | pX459 | | | cacc | |
| 49 | AAACcgctttttgggtactggccc | FrSe2-gRNA_AS | 16743 | gRNA GG oligo | pX459 | | | | aaac |
| 50 | GGGTGAAGACTCcaccgagggcctatttcccatgattcc | U6-gRNA-BbsI_FW | 16744 | PCR primer GG cloning | pTVF-g/go-BsaI | | BbsI | cacc | |
| 51 | GGGTGAAGACGTatgccaaaaaagcaccgactcggtg | U6-gRNA-BbsI_RV | 16745 | PCR primer GG cloning | pTVF-g/go-BsaI | | BbsI | | atgc |
| 52 | GGTGGGTCTCAccgggagggcctatttcccatgattcc | U6-gRNA-BsaI_FW | 16746 | PCR primer GG cloning | pTVF-g/go-BbsI | | BsaI | ccgg | |
| 53 | GGTGGGTCTCGaacccaaaaaagcaccgactcggtg | U6-gRNA-BsaI_FW | 16747 | PCR primer GG cloning | pTVF-g/go-BbsI | | BsaI | | aacc |
| 54 | cgacggccagtgaattcgagctcggtacACAAGTTTGTACAAAAAAGCAGGCTggctccggtgcccgtcagtgg | GAs-E1Fa-mCh_F1_FW | 16748 | PCR primer Gibson Assembly | pUC57-mCherry | attB1 | | | |
| 55 | ggctgtcTCAtacctttctcttctttttggcttgtacagctcgtccatgc | GAs-E1Fa-mCh_F1_RV | 16749 | PCR primer Gibson Assembly | pUC57-mCherry | | | | |

| | | | | | | | | | |
|---|---|---|---|---|---|---|---|---|---|
| 56 | ccaaaaaagaagagaaaggtaTGAgacagcctcgactgtgccttctagttg | GAs-E1Fa-mCh_F2_FW | 16750 | PCR primer Gibson Assembly | pUC57-mCherry | | | | |
| 57 | cagtcgacgggcccgggatccgatatctagACCACTTTGTACAAGAAAGCTGGGTCCATAGAGCCCACCGCATCC | GAs-E1Fa-mCh_F2_RV | 16751 | PCR primer Gibson Assembly | pUC57-mCherry | attB2 | | | |
| 58 | GTGAACACGCTACCGGaCTCGAGAATTCgccac | Mut-BsaI_FW | 16781 | SDM mCherry cassette | pUC57-mCherry | | | | |
| 59 | gtggcGAATTCTCGAGtCCGGTAGCGTGTTCAC | Mut-BsaI_RV | 16782 | SDM mCherry cassette | pUC57-mCherry | | | | |
| 60 | cacccgagaccgagagagggtctcc | pX459-BbsI-to-BsaI_S | 16783 | GG Oligo | pX459 BsaI | | BsaI | cacc | |
| 61 | aaacggagaccctctctcggtctcg | pX459-BbsI-to-BsaI_AS | 16784 | GG Oligo | pX459 BsaI | | BsaI | | aaac |
| 62 | ccgGGGccagtacccaaaaagcGGGGGGGT | FrSe-BsaI_S | 16785 | FrameSelector GG oligo | pTVF | | | ccgg | |
| 63 | aaccACCCCCCCgctttttgggtactggCC | FrSe-BsaI_AS | 16786 | FrameSelector GG oligo | pTVF | | | | aacc |
| 64 | caccCCCCCCCgctttttgggtactggCCC | FrSe-BbsI_S | 16787 | FrameSelector GG oligo | pTVF | | | cacc | |
| 65 | atgcGGGccagtacccaaaaagcGGGGGGG | FrSe-BbsI_AS | 16788 | FrameSelector GG oligo | pTVF | | | | atgc |
| 66 | CATGCGGTGACGTGGAAGAGAATCCCGGCCCTatgattgaacaagatggattgcacgcaggttctcc | GAs-iCG-F1_FW | 16831 | PCR primer Gibson Assembly | iCG | | | | |
| 67 | cactctcggcatggacgagctgtacaagtgaaatcaacctctggattacaaaatttgtg | GAs-iCG-F2_FW | 16832 | PCR primer Gibson Assembly | iCG | | | | |
| 68 | ccagatcttgggtgggttactccagactgccaaggacagggaagggagcagtg | GAs-iCG-F2_RV | 16833 | PCR primer Gibson Assembly | iCG | | | | |
| 69 | ggcagtctggagtaacccacccaagatctgg | GAs-iCG-F3_FW | 16834 | PCR primer Gibson Assembly | iCG | | | | |
| 70 | AGGGCCGGGATTCTCTTCCACGTCAC | GAs-iCG-F3_RV | 16835 | PCR primer Gibson Assembly | iCG | | | | |
| 71 | GGTGGGTCTCAccggGGccagtacccaaaaagcGGGGGGGAGGGACAGCCCCCCCCCAAAG | BsaI-FS-InsL_FW | 16911 | PCR primer cloning | pTFID | | BsaI | ccgg | |
| 72 | GGTGGGTCTCGatgcgGGccagtacccaaaaagcGGGGGGGAGGGACAGCCCCCCCCCAAAG | BsaI-FS-InsR_RV | 16912 | PCR primer cloning | pTFID | | BsaI | | atgc |
| 73 | GGTGGGTCTCGtcctTTTTCCCCGTATCCCCCCAGG | BsaI-DTS-FS-InsL_RV | 17004 | PCR primer cloning | pTFID | | BsaI | | tcct |
| 74 | GGTGGGTCTCGttccTTTTCCCCGTATCCCCCCAGG | BsaI-DTS-FS-InsR_FW | 17005 | PCR primer cloning | pTFID | | BsaI | ttcc | |
| 75 | AGGAatgctttgcatacttctgcctgctggggagcctggggactttccacaccctaactgacacacattccacagc | BsaI-L-DTS_S | 17006 | DTS GG oligo | pTFID | | | agga | |
| 76 | AACCgctgtggaatgtgtgtcagttagggtgtggaaagtccccaggctccccagcaggcagaagtatgcaaagcat | BsaI-L-DTS_AS | 17007 | DTS GG oligo | pTFID | | | | aacc |
| 77 | CACCatgctttgcatacttctgcctgctggggagcctggggactttccacaccctaactgacacacattccacagc | BsaI-R-DTS_S | 17008 | DTS GG oligo | pTFID | | | cacc | |
| 78 | GGAAgctgtggaatgtgtgtcagttagggtgtggaaagtccccaggctccccagcaggcagaagtatgcaaagcat | BsaI-R-DTS_AS | 17009 | DTS GG oligo | pTFID | | | | ggaa |
| 79 | ccggGGCCGGCTTGTCGACGACGGCGGTCTCCGTCGTCAGGATCATCCGG | Bxb1-attBL_S | 17193 | GG Oligo | pTVD-attB (Bxb1) | Bxb1 attB-GT | ccgg | | |
| 80 | aaccCCGGATGATCCTGACGACGGAGACCGCCGTCGTCGACAAGCCGGCC | Bxb1-attBL_AS | 17194 | GG Oligo | pTVD-attB (Bxb1) | Bxb1 attB-GT | | aacc | |
| 81 | caccCCGGATGATCCTGACGACGGAGTCCGCCGTCGTCGACAAGCCGGCC | Bxb1-attBmRev_S | 17195 | GG Oligo | pTVD-attB (Bxb1) | Bxb1 attB-GA | | cacc | |
| 82 | atgcGGCCGGCTTGTCGACGACGGCGGACTCCGTCGTCAGGATCATCCGG | Bxb1-attBmRev_AS | 17196 | GG Oligo | pTVD-attB (Bxb1) | Bxb1 attB-GA | | | atgc |
| 83 | aacgacggccagtgaattcgagctcgGTACACAAGTTTGTACAAAAAAGCAGGCTACCATGGTGAGCAAGGGCGAGGAGC | GA-RGR-F1_FW | 17201 | PCR primer Gibson Assembly | pTVD-ECFP-RGR | | | | |
| 84 | gaaaaacctttacaaccctactgacgaatcTTACTTGTACAGCTCGTCCATGCCG | GA-RGR-F1_RV | 17202 | PCR primer Gibson Assembly | pTVD-ECFP-RGR | | | | |
| 85 | gattcgtcagtagggttgtaaaggtttttcttttcctgagaaaacaaccttttgttttctcaggttttgctttttggcctttccctagctttaaaaaaaaaaaagcaaaa | GA-RGR-MALAT1_S | 17203 | Oligo Gibson Assembly | pTVD-ECFP-RGR | | | | |
| 86 | cctttccctagctttaaaaaaaaaaaagcaaaagaaacaccgggtcttcgagaagac | GA-RGR-F2_FW | 17204 | PCR primer Gibson Assembly | pTVD-ECFP-RGR | | | | |
| 87 | cgccatgccgaagcatgttgcccagccggcgccagcgaggaggctgggaccatgccggcccaaaaaagcaccgactcggtgc | GA-RGR-F2_RV | 17205 | PCR primer Gibson Assembly | pTVD-ECFP-RGR | | | | |
| 88 | gccggctgggcaacatgcttcggcatggcgaatgggacgaattctaactagagctcgctgatcagc | GA-RGR-F3_FW | 17206 | PCR primer Gibson Assembly | pTVD-ECFP-RGR | | | | |
| 89 | actgccttgggaaaagcgcctcccctaccctccccagcatgcctgctattctc | GA-RGR-F3_RV | 17207 | PCR primer Gibson Assembly | pTVD-ECFP-RGR | | | | |
| 90 | gggtaggggaggcgcttttcc | GA-RGR-F4_FW | 17208 | PCR primer Gibson Assembly | pTVD-ECFP-RGR | | | | |
| 91 | cagtcgacgggcccgggatccgatatctagACCACTTTGTACAAGAAAGCTGGGTcgaaaggcccggagatgagg | GA-RGR-F4_RV | 17209 | PCR primer Gibson Assembly | pTVD-ECFP-RGR | | | | |
| 92 | tgtacaagtaaagcggccgcgactCTAG | GA-TCVLP-F1_FW | 17220 | PCR primer Gibson Assembly | pTVF-ID-gRNA-LP | | | | |
| 93 | ccaggcgatctgacggttcactaaacgagc | GA-TCVLP-F1_RV | 17221 | PCR primer Gibson Assembly | pTVF-ID-gRNA-LP | | | | |
| 94 | gctcgtttagtgaaccgtcagatcgcctggccggacgtcttcgagaagaccgggttCAAG | GA-TCVLP-F2_FW | 17222 | PCR primer Gibson Assembly | pTVF-ID-gRNA-LP | | | | |
| 95 | cgatatctagAaatgcaggtcttctcgaagaccc | GA-TCVLP-F2_RV | 17223 | PCR primer Gibson Assembly | pTVF-ID-gRNA-LP | | | | |
| 96 | cttcgagaagacctgcattTctagatatcgATGGCCAAGTTGACCAGTGCCG | GA-TCVLP-F3_FW | 17224 | PCR primer Gibson Assembly | pTVF-ID-gRNA-LP | | | | |
| 97 | tTCTAGagtcgcggccgctttacttgtacaTCAGTCCTGCTCCTCGGCCACG | GA-TCVLP-F3_RV | 17225 | PCR primer Gibson Assembly | pTVF-ID-gRNA-LP | | | | |
| 98 | GGGGACAAGTTTGTACAAAAAAGCAGGCTAAGGATCTGCGATCGCTCCGGTGC | attB1-E1Fa_FW | 17226 | PCR primer Gateway cloning | pENTR-GFP-puro | attB1 | | | |
| 99 | GGGGACCACTTTGTACAAGAAAGCTGGGTgtatgctatacgaagttatagatctAGGGACAGC | attB2-Ins_RV | 17227 | PCR primer Gateway cloning | pENTR-GFP-puro | attB2 | | | |
| 100 | ccggGTGGTTTGTCTGGTCAACCACCGCGGTCTCAGTGGTGTACGGTACAAACCC | Bxb1-attPL_S | 17228 | GG Oligo | pTVF-ID-attP (Bxb1) | Bxb1 attP-GT | ccgg | | |
| 101 | aaccGGGTTTGTACCGTACACCACTGAGACCGCGGTGGTTGACCAGACAAACCAC | Bxb1-attPL_AS | 17229 | GG Oligo | pTVF-ID-attP (Bxb1) | Bxb1 attP-GT | | aacc | |
| 102 | caccGGGTTTGTACCGTACACCACTGAGTCCGCGGTGGTTGACCAGACAAACCAC | Bxb1-attPmRev_S | 17230 | GG Oligo | pTVF-ID-attP (Bxb1) | Bxb1 attP-GA | | cacc | |
| 103 | atgcGTGGTTTGTCTGGTCAACCACCGCGGACTCAGTGGTGTACGGTACAAACCC | Bxb1-attPmRev_AS | 17231 | GG Oligo | pTVF-ID-attP (Bxb1) | Bxb1 attP-GA | | | atgc |
| 104 | caccGAACCGATCGTCTCGTGGCCTGTCTGATGAGTCCGTGAGGACGAAACGAGTAAGCTCGTCACAGGCACTGCACCCGTTCG | RGR-gRNA-STK11L_S | 17314 | GG Oligo | pTVD-ECFP-RGR-STK11L | | | | |
| 105 | aaacCGAACGGGTGCAGTGCCTGTGACGAGCTTACTCGTTTCGTCCTCACGGACTCATCAGACAGGCCACGAGACGATCGGTTC | RGR-gRNA-STK11L_AS | 17315 | GG Oligo | pTVD-ECFP-RGR-STK11L | | | | |
| 106 | caccGAACCGATCGTCTCGTGCGGCCTCTGATGAGTCCGTGAGGACGAAACGAGTAAGCTCGTCAGGCCGTTGGCAATCTCGGG | RGR-gRNA-STK11R2_S | 17316 | GG Oligo | pTVD-ECFP-RGR-STK11R2 | | | | |
| 107 | aaacCCCGAGATTGCCAACGGCCTGACGAGCTTACTCGTTTCGTCCTCACGGACTCATCAGAGGCCGCACGAGACGATCGGTTC | RGR-gRNA-STK11R2_AS | 17317 | GG Oligo | pTVD-ECFP-RGR-STK11R2 | | | | |
| 108 | caccGAACCGATCGTCTCGTGCGCCACCTGATGAGTCCGTGAGGACGAAACGAGTAAGCTCGTCGTGGCGGGGTCGGCGCAGTT | RGR-gRNA-CDKN2AL_S | 17318 | GG Oligo | pTVD-ECFP-RGR-CDKN2AL | | | | |
| 109 | aaacAACTGCGCCGACCCCGCCACGACGAGCTTACTCGTTTCGTCCTCACGGACTCATCAGGTGGCGCACGAGACGATCGGTTC | RGR-gRNA-CDKN2AL_AS | 17319 | GG Oligo | pTVD-ECFP-RGR-CDKN2AL | | | | |
| 110 | caccGAACCGATCGTCTCGTGCGGGTCCTGATGAGTCCGTGAGGACGAAACGAGTAAGCTCGTCGACCCGTGCACGACGCTGCC | RGR-gRNA-CDKN2AR_S | 17320 | GG Oligo | pTVD-ECFP-RGR-CDKN2AR | | | | |
| 111 | aaacGGCAGCGTCGTGCACGGGTCGACGAGCTTACTCGTTTCGTCCTCACGGACTCATCAGGACCCGCACGAGACGATCGGTTC | RGR-gRNA-CDKN2AR_AS | 17321 | GG Oligo | pTVD-ECFP-RGR-CDKN2AR | | | | |
| 112 | GGTGgaagacctcaccGAACCGATCGTCTCGTG | RGRcas-gRNA-BbsI-L_FW | 17322 | PCR primer GG cloning | RGR-gRNA donor | | BbsI | cacc | |
| 113 | GGTGgaagacctACTCctgatcagcgagctctagttagaattcg | RGRcas-gRNA-BbsI-R1_RV | 17323 | PCR primer GG cloning | RGR-gRNA donor | | BbsI | | actc |
| 114 | GGTGgaagacctGAGTGAACCGATCGTCTCGTG | RGRcas-gRNA-BbsI-L1_FW | 17324 | PCR primer GG cloning | RGR-gRNA donor | | BbsI | gagt | |
| 115 | GGTGgaagacctaaacctgatcagcgagctctagttagaattcg | RGRcas-gRNA-BbsI-R_RV | 17325 | PCR primer GG cloning | RGR-gRNA donor | | BbsI | | aaac |

| | | | | | | | | | |
|---|---|---|---|---|---|---|---|---|---|
| 116 | GGTGgaagacctCGGCctgatcagcgagctctagttagaattcg | RGRcas-gRNA-BbsI-R2_RV | 17326 | PCR primer GG cloning | RGR-gRNA donor | | BbsI | | cggc |
| 117 | GGTGgaagacctGCCGGAACCGATCGTCTCGTG | RGRcas-gRNA-BbsI-L2_FW | 17327 | PCR primer GG cloning | RGR-gRNA donor | | BbsI | gccg | |
| 118 | GGTGgaagacctAGCCctgatcagcgagctctagttagaattcg | RGRcas-gRNA-BbsI-R3_RV | 17328 | PCR primer GG cloning | RGR-gRNA donor | | BbsI | | agcc |
| 119 | GGTGgaagacctGGCTGAACCGATCGTCTCGTG | RGRcas-gRNA-BbsI-L3_FW | 17329 | PCR primer GG cloning | RGR-gRNA donor | | BbsI | ggct | |
| 120 | ccggTATCGCAGGTGCACCTGCATCT | LBbsI-to-PaqCI_S | 17538 | BsaI to PaqCI GG oligo | pTCV-PaqCI | | PaqCI | ccgg | aacc |
| 121 | aaccAGATGCAGGTGCACCTGCGATA | LBbsI-to-PaqCI_AS | 17539 | BsaI to PaqCI GG oligo | pTCV-PaqCI | | PaqCI | ccgg | aacc |
| 122 | caccTATCGCAGGTGCACCTGCATCT | RBbsI-to-PaqCI_S | 17540 | BbsI to PaqCI GG oligo | pTCV-PaqCI | | PaqCI | cacc | atgc |
| 123 | atgcAGATGCAGGTGCACCTGCGATA | RBbsI-to-PaqCI_AS | 17541 | BbsI to PaqCI GG oligo | pTCV-PaqCI | | PaqCI | cacc | atgc |
| 124 | GGTGGGTCTCAccggACgtcttcgagaagacCGggttccagtgaattcgagctcgGTACCccg | BsaI-BbsI-FS-InsL_FW | 17825 | PCR primer GG cloning | pTVF-ID-BbsI-PaqCI | | BsaI | ccgg | |
| 125 | GGTGggtctcgaaccAGATGCAGGTGCACCTGCGATAccggCGTTCAGCTTTTTTGTACAAACTTGaacc | BsaI-InsL-PaqCI_RV | 17826 | PCR primer GG cloning | pTVF-ID-BbsI-PaqCI | | BsaI | | aacc |
| 126 | GGTGGGTCTCAcaccTATCGCAGGTGCACCTGCATCTgcatGTGATCGATTCGACAGATCacacc | BsaI-PaqCI-InsR_FW | 17827 | PCR primer GG cloning | pTVF-ID-BbsI-PaqCI | | BsaI | cacc | |
| 127 | GGTGggtctcgatgcAGgtcttctcgaagacCCggtggcccgggatccgatatctagAaatgc | BsaI-BbsI-FS-InsR_RV | 17828 | PCR primer GG cloning | pTVF-ID-BbsI-PaqCI | | BsaI | | atgc |
| 128 | ccggaaataatgattttattttgactgatagtgacctgttcgttgcaacaaattgataagcaatgcttttttataatgccaactttgtatagaaaagttgCTGGCA | GG-attL4-BC1_S | 18304 | Golden Gate oligonucleotide; MultiSite Gat | pTCV-attL4-R1-R2-attL3 | | | | |
| 129 | cgttTGCCAGcaacttttctatacaaagttggcattataaaaaagcattgcttatcaatttgttgcaacgaacaggtcactatcagtcaaaataaaatcattattt | GG-attL4-BC1_AS | 18305 | Golden Gate oligonucleotide; MultiSite Gat | pTCV-attL4-R1-R2-attL3 | | | | |
| 130 | GCTGACTGTGcaactttattatacaaagttggcattataaaaaagcattgcttatcaatttgttgcaacgaacaggtcactatcagtcaaaataaaatcattattt | GG-attL3-BC3_S | 18322 | Golden Gate oligonucleotide; MultiSite Gat | pTCV-attL4-R1-R2-attL3 | | | | |
| 131 | ATGCaaataatgattttattttgactgatagtgacctgttcgttgcaacaaattgataagcaatgcttttttataatgccaactttgtataataaagttgCACAGT | GG-attL3-BC3_AS | 18323 | Golden Gate oligonucleotide; MultiSite Gat | pTCV-attL4-R1-R2-attL3 | | | | |
| 132 | ccagtgaattcgagctcgGTACCccggTATCGCAGGTGCACCTGCATCTggtttcaactttgtatagaaaagttgaac | GA-PaqCI-attR4_FW | 18324 | PCR primer Gibson Assembly | pTCV-attR4-attR3-PaqCI | | | | |
| 133 | cgggcccgggatccgatatctagAaatgcAGATGCAGGTGCACCTGCGATAggtgcaactatgtataataaagttgaacgag | GA-PaqCI-attR3_RV | 18325 | PCR primer Gibson Assembly | pTCV-attR4-attR3-PaqCI | | | | |
| 134 | cctctgacacatgcagctcccgg | GA-pTCVKan_FW | 18789 | PCR primer Gibson Assembly | pTCVKan-attR4-attR3-PaqCI | | | | |
| 135 | ccgggagctgcatgtgtcagaggCCGGAATTGCCAGCTGGGGC | GA-Kan_FW | 18790 | PCR primer Gibson Assembly | pTCVKan-attR4-attR3-PaqCI | | | | |
| 136 | GGAGGGGGTTCCTTCTCTG | TP53-Ex2-3_for | 732 | PCR TP53 exon 3 | | | | | |
| 137 | GGACAAACCACAACTAGAATGCA | RIP-S | 870 | PCR detection of iCGC, NeoR | | | | | |
| 138 | AAGCCCAGACGGAAACCG | p53-E4R | 1399 | PCR TP53 exon 3 | | | | | |
| 139 | GCTCCTGAGGTGTAGACGCCAA | T7_p53 Exon 5_fw | 6557 | PCR TP53 Exon 5 | | | | | |
| 140 | GCACATCTCATGGGGTTATAGGGA | T7_p53 Exon 5_rev | 6558 | PCR TP53 Exon 5 | | | | | |
| 141 | GAATGGCCTGTTCGGAAACCT | Cas9-For | 8240 | PCR detection of iCGC, Cas9 | | | | | |
| 142 | CATCCGCTCGATGAAGCTCT | Cas9-Rev | 8241 | PCR detection of iCGC, Cas9 | | | | | |
| 143 | GAGCAGCGAGAAGCGTTCAGAG | HR700PA_1_Core_Isul2_For | 8254 | PCR pTVF integration | | | | | |
| 144 | TCCAGGGGTCCGAGAGCTCAGCTAG | prAAVS1_S | 9451 | PCR pTVF integration into AAVS1, left | | | | | |
| 145 | tttctggattcatcgactgtgg | Puro 4-s | 11072 | PCR iCGC integration | | | | | |
| 146 | GGCTCCGATCAGGTTCTTC | LSL-Cas9-loxP-Cas9-rev2 | 11924 | PCR iCGC integration | | | | | |
| 147 | TGCGACGTTGCTGTCAAAAG | hSH231_PCR-intL_FW1 | 15013 | PCR iCGC integration in SHS231, left | | | | | |
| 148 | CCACTCATTAAGCACAAAAGATACT | hSH231_PCR-intR_RV1 | 15018 | PCR iCGC integration in SHS231, right | | | | | |
| 149 | CGGACCGCTATCAGGACATA | hSH231_PCR-intR_FW2 | 15019 | PCR detection of iCGC, NeoR | | | | | |
| 150 | ccagtgaattcgagctcggtacGAGGCTGGCCAAAGGAATAAG | GAs-SH231L_FW | 15658 | PCR iCGC integration in SHS231 | | | | | |
| 151 | ctcatcaatgtatcttaGCAAAATTTTCTATCTTCTTAGTGG | GAs-SH231R_FW | 15662 | PCR iCGC integration in SHS231 | | | | | |
| 152 | GGGgaagactgatgcGAGCAGAGCCAGGAACCCCTG | BbsI-AAVS1_RV | 15667 | PCR pTVF integration into AAVS1, right | | | | | |
| 153 | caccGTCGAGTCGCTTCTCGATTA | gRNA1-hR26_S | 15678 | PCR pTVF integration into ROSA26, left | | | | | |
| 154 | CCCTTTGCTATTTTGCCCGT | G-CDKN2A_FW | 15704 | PCR CDKN2A Exon 2 | | | | | |
| 155 | GCTCTGAGCTTTGGAAGCTCTC | T7-CDKN2A_RV | 15706 | PCR CDKN2A Exon 2 | | | | | |
| 156 | TGTGCACAAGGACATCAAGC | G-STK11_FW | 15707 | PCR STK11 exon 5 | | | | | |
| 157 | CTCTGTGAAGATCCCTGACGG | G-STK11_RV | 15708 | PCR STK11 exon 5 | | | | | |
| 158 | GGAggtctcTccggCATTCTGCGCTTGGATATACAGC | BsaI-CDKN2A_FW | 15715 | PCR CDKN2A Exon 2 | | | | | |
| 159 | GGGgaagactgatgcGGCCTTGAACTAGCAGAGGG | BbsI-CDKN2A_RV | 15718 | PCR CDKN2A Exon 2 | | | | | |
| 160 | GGAggtctcTccggCCTGTGGACATCCGGGGCCC | BsaI-STK11_FW | 15719 | PCR STK11 exon 5 | | | | | |
| 161 | GGGgaagactgatgcCACCTTCCACTAGTGGAGGC | BbsI-STK11_RV | 15722 | PCR STK11 exon 5 | | | | | |
| 162 | GGAggtctcTccggGACACCTGATTGGCTGTCGAAGC | BsaI-hR26_FW | 15727 | PCR iCGC integration in ROSA26 | | | | | |
| 163 | GGGgaagactgatgcGGTATAAAACTCAGGTGATCATGC | BbsI-hR26_RV | 15728 | PCR iCGC integration in ROSA26 | | | | | |
| 164 | gaagactgatgcCCTTCAGACTCAGTGAGCACATtaagatacattgatgagtttggacaaaccacaactagaatg | iCGC-HITI1_RV | 15768 | PCR detection of iCGC, NeoR | | | | | |
| 165 | cgacggccagtgaattcgagctcggtacacaagtttgtacaaaaaagcaggctgaggatcctttaccactccc | GAs-iCGC-F1_FW | 16676 | PCR iCGC integration | | | | | |
| 166 | cgacggccagtgaattcgagctcggtacacaagtttgtacaaaaaagcaggctgaggatcctttaccactccc | GAs-iCGC-F1_FW | 16676 | PCR detection of iCGC, Cas9 | | | | | |
| 167 | tcctcgcccttgctcaccat | GAs-iCGC-F2_RV | 16678 | PCR landing pad detection (EF1a) | | | | | |
| 168 | agacgagtcggatctcccttt | SPLINTR-SEQ-FW | 16692 | PCR landing pad detection (poly A) | | | | | |
| 169 | AGGAatgctttgcatacttctgcctgctggggagcctggggactttccacaccctaactgacacacattccacagc | BsaI-L-DTS_S | 17006 | PCR landing pad detection (poly A) | | | | | |
| 170 | AACCgctgtggaatgtgtgtcagttagggtgtggaaagtccccaggctccccagcaggcagaagtatgcaaagcat | BsaI-L-DTS_AS | 17007 | PCR landing pad integration (internal) | | | | | |
| 171 | CACCatgctttgcatacttctgcctgctggggagcctggggactttccacaccctaactgacacacattccacagc | BsaI-R-DTS_S | 17008 | PCR landing pad integration (internal) | | | | | |
| 172 | GGAAgctgtggaatgtgtgtcagttagggtgtggaaagtccccaggctccccagcaggcagaagtatgcaaagcat | BsaI-R-DTS_AS | 17009 | PCR landing pad detection (EF1a) | | | | | |
| 173 | gggtaggggaggcgcttttcc | GA-RGR-F4_FW | 17208 | PCR gRNA-LP detection and integration | | | | | |
| 174 | tTCTAGagtcgcggccgctttacttgtacaTCAGTCCTGCTCCTCGGCCACG | GA-TCVLP-F3_RV | 17225 | PCR gRNA-LP detection and integration | | | | | |
| 175 | GAACTCCACGCCGTTCAGGGTG | copGFP_rev1 | 19160 | PCR gRNA-LP detection and integration | | | | | |
